# Stem cell heterogeneity and reiteration of developmental signaling underlie melanocyte regeneration in zebrafish

**DOI:** 10.1101/2022.04.27.489712

**Authors:** William Tyler Frantz, Sharanya Iyengar, James Neiswender, Alyssa Cousineau, Rene Maehr, Craig J. Ceol

**Author notes:** Corresponding author: Craig J. Ceol.

## Abstract

Tissue-resident stem cells are present in many adult organs, where they are important for organ homeostasis and repair in response to injury. However, the signals that activate these cells and the mechanisms governing how these cells self-renew or differentiate are highly context-dependent and incompletely understood, particularly in non-hematopoietic tissues. In the skin, melanocyte stem cells (McSCs) are responsible for replenishing mature pigmented melanocytes. In mammals, these cells reside in the hair follicle bulge and bulb niches where they are activated during homeostatic hair follicle turnover and following melanocyte destruction, as occurs in vitiligo and other skin hypopigmentation disorders. Recently, we identified adult McSCs in the zebrafish. To elucidate mechanisms governing McSC self-renewal and differentiation we analyzed individual transcriptomes from thousands of melanocyte lineage cells during the regeneration process. We identified transcriptional signatures for McSCs, deciphered transcriptional changes and intermediate cell states during regeneration, and analyzed cell-cell signaling changes to discover mechanisms governing melanocyte regeneration. We identified KIT signaling via the RAS/MAPK pathway as a regulator of McSC direct differentiation. Analysis of the scRNAseq dataset also revealed a population of *mitfa*/*aox5* co-expressing cells that divides following melanocyte destruction, likely corresponding to cells that undergo self-renewal. Our findings show how activation of different subpopulations of *mitfa*-positive cells underlies self-renewal and differentiation to properly reconstitute the melanocyte pigmentary system following injury.

## Introduction

Adult stem cells are responsible for adult tissue maintenance and critical in recovery from injury. These cells are maintained through self-renewal and respond to tissue injury by proliferating or differentiating. Melanocyte stem cells (McSCs), which are found in the hair follicle niche, are adult stem cells that replenish melanocytes, which ultimately impart pigment to hair and skin (Nishimura et al., 2002; Slominski et al., 2005). Mechanisms governing adult stem cell behavior are critical to maintaining proper tissue function as well as recovery from injury or disease.

Zebrafish, with their characteristic stripes of pigment-retaining dermal melanocytes, have emerged as a useful model to study melanocyte regeneration. Ablation of embryonic and adult melanocytes results in generation of new melanocytes from tissue-resident stem cells (Hultman et al., 2009; O’Reilly-Pol and Johnson, 2013; Rawls and Johnson, 2000; Yang et al., 2007). These tissue-resident stem cells were identified in adult zebrafish stripes as unpigmented cells that are admixed with mature melanocytes and express *mitfa*, the zebrafish ortholog of the MITF melanocyte lineage regulator (Iyengar et al., 2015). These zebrafish McSCs respond to injury by either differentiating into mature melanocytes to reconstitute the skin’s pigment pattern or dividing symmetrically or, less commonly, asymmetrically to replenish the stem cell pool. Daughter cells from these divisions can, following subsequent injury, directly differentiate, indicating a plasticity of stem cell fates that can be adopted by McSCs (Iyengar *et al*., 2015). Other tissue-resident adult stem cells, such as murine basal epidermal stem cells, can also adopt different fates during regeneration. These epidermal stem cells appear initially uncommitted, with environmental signals likely to trigger self-renewal or differentiation (Rompolas et al., 2016).

Identification of zebrafish McSCs facilitated our investigation of mechanisms governing differentiation. *In vivo* imaging revealed that the unpigmented McSCs upregulate WNT signaling preceding undergoing differentiation, and animals treated with WNT inhibitor fail to regenerate. Meanwhile, self-renewal is unaffected, demonstrating a fate-specific requirement for WNT-signaling (Iyengar *et al*., 2015). Like in the zebrafish, previous murine studies have identified coordinated WNT signaling as a key regulator of differentiation during melanocyte regeneration (Rabbani et al., 2011). More recently, profiling of McSC and melanocyte single cell transcriptomes from regenerating hair follicles has revealed coordinated activation of WNT/BMP signaling as a gate governing differentiation (Infarinato et al., 2020). These studies underline the conserved mechanisms underpinning melanocyte regeneration in murine and zebrafish models, emphasizing the utility of studying adult McSCs in zebrafish.

While the mechanisms governing stem cell fates during regeneration have been elucidated in some tissues, adult stem cell identities and responses to injury are incompletely understood. Some tissue-resident stem cells, such as epidermal hair follicle and basal epidermal stem cells, rely on positional cues for fate determination, suggestive of a stochastic model (Rompolas and Greco, 2014; Rompolas *et al*., 2016). Yet, other tissue resident stem cells, such as hematopoietic and lung epithelial stem cells, appear to be organized in a hierarchical manner with a self-renewing common multipotent stem cell giving rise to lineage-committed progeny (Laurenti and Göttgens, 2018; McQualter et al., 2010). Even within the same stem cell pool the behaviors can be context dependent. In response to injury, murine McSCs will prioritize differentiation over self-renewal, but in normal hair follicle turnover these cells engage in balanced differentiation and self-renewal (Chou et al., 2013; Nishimura *et al*., 2002). It is currently unknown if the different McSC behaviors observed in murine and zebrafish regeneration studies are due to stem-cell heterogeneity like lung stem cells or are stochastically regulated like epidermal stem cells. To date, most analysis of McSCs have relied on *in vivo* imaging and targeted transcriptomics, potentially missing the role of mixed transcriptional responses to injury. Single cell RNA sequencing has been used to dissect the extraordinary heterogeneity and transcription dynamics present in developing pigment cell lineages as well as melanocyte regeneration and offers new powerful insights into signaling mechanisms governing McSC fate (Infarinato *et al*., 2020; Saunders et al., 2019).

Here, we utilize single cell transcriptomics to find that zebrafish McSCs are a heterogenous group of *mitfa*-expressing cells within the melanocyte stripe which require KIT signaling to directly differentiate during melanocyte regeneration. Importantly, changes in gene expression of McSCs during regeneration are well conserved between mice and zebrafish. Lastly, we observe that heterogeneity of McSCs, evident in distinct transcriptional signatures, underlies the different fates adopted by these cells – differentiation versus self-renewal – during regeneration.

## Results

### Adult melanocyte lineage cells revealed by single-cell RNA-sequencing

During melanocyte regeneration, unpigmented *mitfa*-expressing cells in the zebrafish stripe engage in coupled differentiation and self-renewal to replenish lost melanocytes and maintain the stem cell pool (Iyengar *et al*., 2015). To identify the mechanisms governing these behaviors we sought to capture cell states and transcriptional changes in the melanocyte lineage during regeneration. McSCs are rare, so we generated an *Tg(mitfa:nlsEGFP)* reporter line and sorted EGFP-positive cells to isolate the 0.19% of total cells which express *mitfa* (Figure 1A and Supplemental Figure 1A). We utilized neocuproine to ablate mature melanocytes, leading to the destruction of stripe melanocytes (O’Reilly-Pol and Johnson, 2008), then performed scRNAseq on *mitfa:nlsEGFP*-positive cells following melanocyte destruction (Figure 1B). In all, we obtained transcriptomes of 29,453 wild-type cells across six time points before, during, and after melanocyte regeneration. Quality control and pre-processing followed by dimensionality reduction and unsupervised clustering identified several subpopulations of cells (Figure 1C). Analysis of captured cells revealed that 87.5% were positive for endogenous *mitfa* expression, reinforcing the success of our enrichment strategy (Supplemental Figure 1B). The vast majority of *mitfa*-expressing cells were in the two large subgroups, and expression of *sox10*, *tryp1b*, *aox5* and other markers was consistent with these subgroups containing pigment cells (Figure 1C, D). Expression of the *tryp1b* melanin biosynthesis gene was limited to one subpopulation, and we have assigned this subpopulation as differentiated (or differentiating) melanocytes. Expression of another differentiated melanocyte marker, *pmela*, confirmed this assignment (Figure 1E). The two large subgroups were distinguished from one another by their differential expression of the *aox5* gene (Figure 1D). *aox5* is expressed in differentiated xanthophores, a pteridine-producing pigment cell type of zebrafish, and in some undifferentiated pigment progenitor cells (McMenamin et al., 2014; Parichy et al., 2000; Saunders *et al*., 2019). The non-*mitfa* expressing clusters comprised other cell populations such as iridophores, keratinocytes, fibroblasts, immune cells, and Schwann cells, based on expression of canonical cell-type marker genes (Figure 1C,E, Supplemental Figure 1C,D). Our cell-type assignments generally agree with the assignments from a *sox10*-enriched transcriptomic analysis of zebrafish pigment cells in development, supporting the robustness of our approach to capture these cell types (Supplemental Figure 1E) (Saunders *et al*., 2019).

**Figure 1.**
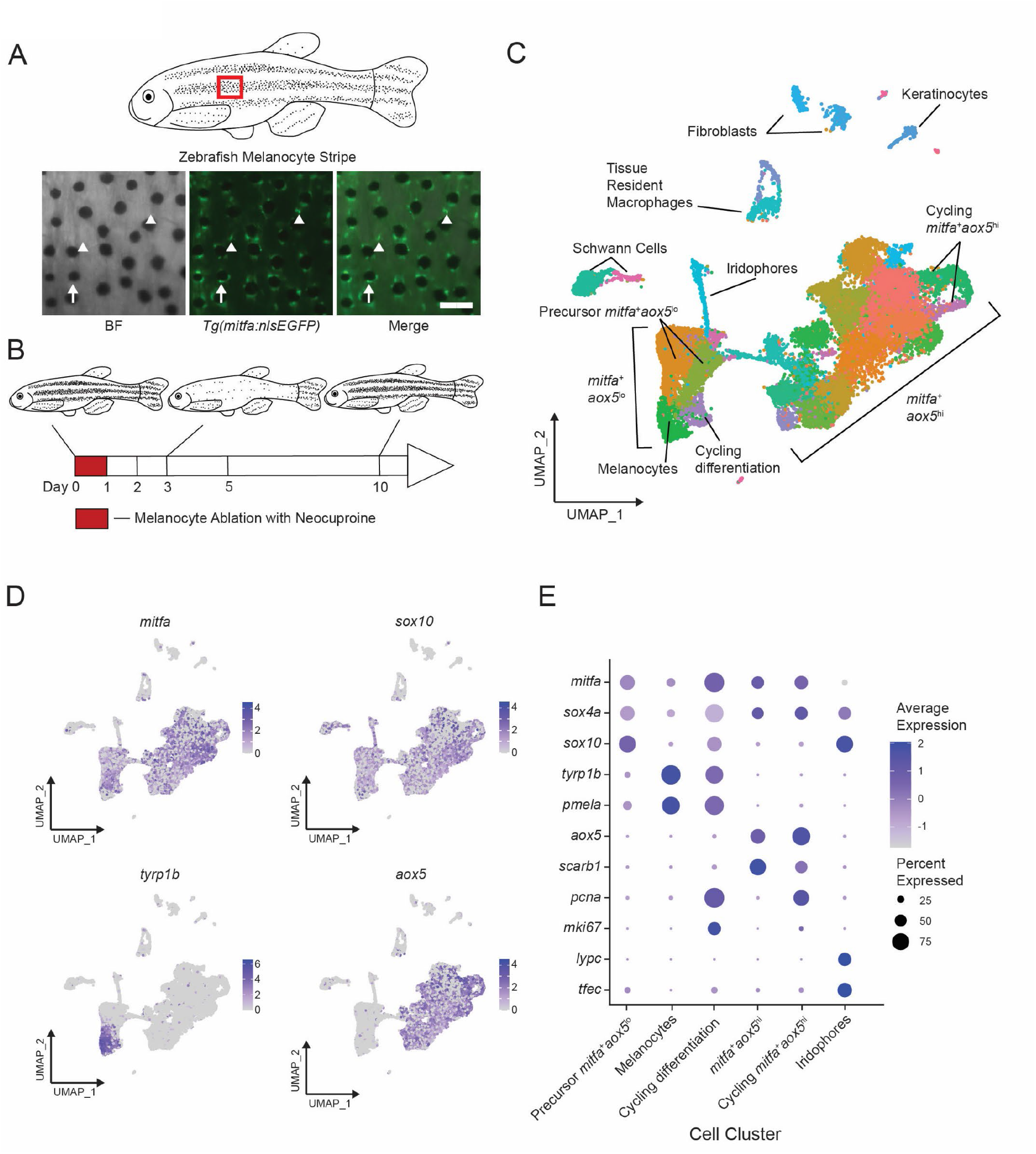
Single-cell transcriptomic identification of melanocyte lineage cells during regeneration. (A) Top, diagram of zebrafish flank with melanocyte stripes. Bottom, representative images of the melanocyte stripe in a *Tg(mitfa:nlsEGFP*) zebrafish. Melanocyte stem cells (McSCs) are unpigmented GFP-expressing cells (arrowheads) admixed with pigmented melanocytes (arrows). Animals were treated with epinephrine prior to imaging to concentrate melanosomes into the cell body of melanocytes. Scale bar = 100µm. (B) Experimental design for transcriptional profiling of McSCs in *Tg(mitfa:nlsEGFP)* zebrafish during melanocyte ablation and regeneration. Cells from *Tg(mitfa:nlsEGFP)* zebrafish were sampled at the days specified. (C) UMAP of cell-type assignments for clusters of cells obtained from *Tg(mitfa:nls:EGFP)* zebrafish. Cells from all time points were included. Coloring is according to unsupervised clustering (Blondel et al., 2008; Stuart et al., 2019). Cells are labeled based on gene expression patterns revealed in panels (D) and (E) and Supplemental Figure 1D. The large group of cells to the bottom left in the UMAP are *mitfa*^+^*aox5*^lo^. Within this group of cells, the *mitfa*^+^*aox5*^lo^*tyrp1b*^+^*pcna^-^* cells are designated as melanocytes, the *mitfa*^+^*aox5*^lo^*tyrp1b*^+^*pcna*^+^ are designated ‘cycling differentiation’, and the other two populations are designated ‘precursor *mitfa*^+^*aox5*^lo^’. The larger group of cells to the bottom right in the UMAP are *mitfa*^+^*aox5*^hi^. Within this group of cells two clusters are *pcna^+^* and are designated ‘cycling *mitfa*^+^*aox5*^hi^’ (n=29,453 cells). (D) Expression of pigment cell markers *mitfa* and *aox5*, melanin biosynthesis gene *tyrp1b* and pigment cell progenitor marker *sox10* shown as feature plots on the UMAP plot from (C). (E) Expression of pigment cell markers, the stem cell gene *sox4a* and cell cycle markers for cell clusters shown in panel C. Dot sizes represent percentage of cells in the cluster expressing the marker and coloring represents average expression.

### Single-cell transcriptomics reveal McSC heterogeneity and dynamic gene expression changes following melanocyte destruction

To investigate if McSCs were clustered in the same *mitfa*^+^*aox5*^lo^ subgroup as melanocytes, we sought to understand the transcriptional and population changes of this subgroup during regeneration. Unsupervised clustering of the *mitfa*^+^*aox5*^lo^ subgroup from integrated samples from the six time points before, during, and after melanocyte regeneration revealed four distinct subpopulations in addition to differentiated melanocytes (Figure 2A). Two of these subpopulations were characterized by low levels of *tryp1b* expression and high levels of *sox4a* (Figure 2B, Supplemental Figure 2A). SOX4 in mammalian cells has been shown in both normal and tumor cells to maintain an undifferentiated state that is associated with stemness (Uy et al., 2015; Vervoort et al., 2013). Because of this as well as the expression of genes typically observed in the neural crest (Supplemental Figure 2A,B), the tissue from which melanocytes are derived during embryonic development, we considered these two subpopulations as putative McSCs and have designated them as McSC-0 and McSC-1. The McSC-0 and McSC-1 subpopulations themselves showed gene expression differences (e.g., *dio3a* and *bco2b*) that drove their separate cluster assignments (Supplemental Figure 2A,C). The remaining three subpopulations expressed moderate to high levels of *tryp1b* (Figure 2B). The subpopulation with highest *tryp1b* was assigned as mature melanocytes, whereas the two subpopulations with moderate *tryp1b* levels also expressed neural crest markers (Supplemental Figure 2A), suggesting that they were not fully differentiated. To validate that our scRNAseq procedure was reproducible enough to enable comparisons between samples, we performed independent sampling of *mitfa*-positive cells without any melanocyte ablation. The cell sampling prevalence between single-cell runs was consistent, as seen by comparing proportions of cell types from two independent, pre-ablation samples (Supplemental Figure 2D), giving confidence to our inter-sample comparisons. Comparisons of samples across different time points shows dynamic shifts in subpopulations following ablation and during regeneration (Figure 2C,D). As expected, the melanocyte subpopulation decreased substantially (by >95%) following neocuproine-mediated melanocyte destruction between day 0 and day 1. The mature melanocyte subpopulation then increased in size as regeneration proceeded through day 10. Comparisons of subpopulations across regeneration also revealed changes in the two subpopulations that express neural crest markers and moderate levels of *tryp1b* (Supplemental Figure 2A,B,C). One of the intermediate subpopulations expressed high levels of cell-cycle genes, including *pcna* and *cdk1* (Figure 2B, Supplemental Figure 2C). We termed this subpopulation “cycling differentiation”. The other intermediate population was also positionally between *tyrp1b*-negative and *tyrp1b*-positive cells but did not express cell-cycle genes (Figure 2B, Supplemental Figure 2C), so we termed this subpopulation “direct differentiation”. Both the cycling differentiation and direct differentiation subpopulations grew following ablation, but then diminished as regeneration neared completion at day 10 (Figure 2C,D).

**Figure 2.**
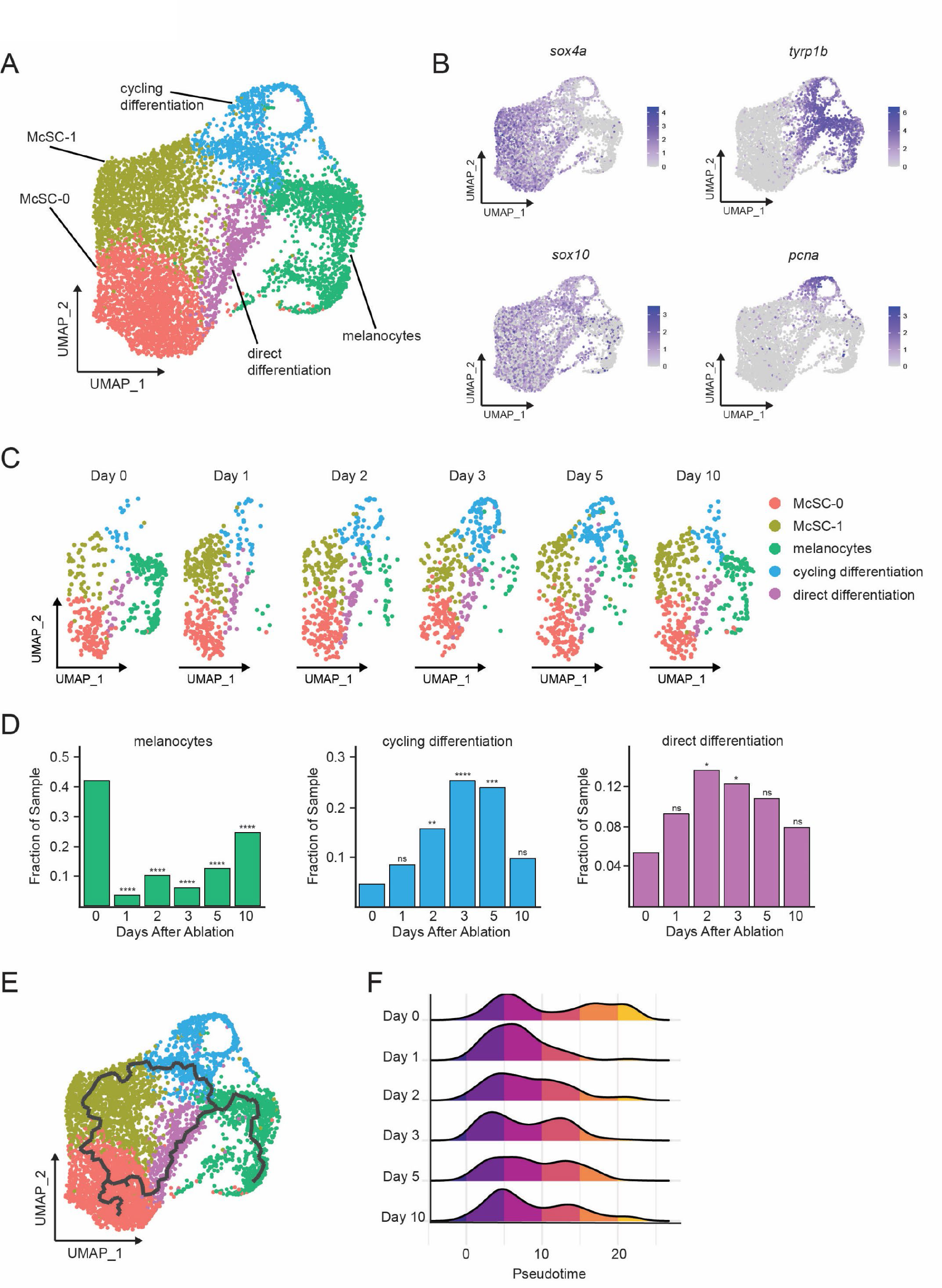
Single-cell transcriptomics reveal dynamic cell and gene expression changes following melanocyte ablation. (A) Integrated sub-clustering of *mitfa^+^aox5*^lo^ cells before, during, and after melanocyte regeneration. (n=5,619 cells). Coloring is according to unsupervised clustering (Blondel *et al*., 2008; Stuart *et al*., 2019). Cell clusters are labeled based on gene expression patterns and inferred trajectories during regeneration, as revealed in panels 2B-F and Supplemental Figure 2. Melanocyte and cycling differentiation clusters are the same as in Figure 1A, whereas the two precursor *mitfa*^+^*aox5*^lo^ clusters from Figure 1A are now resolved into three clusters: McSC-0, McSC-1 and ‘direct differentiation’, the latter of which expresses the stem cell marker *sox4a*, the melanin biosynthesis gene *tyrp1b* and is *pcna*-negative. (B) Expression of stem cell marker *sox4a*, melanin biosynthesis gene *tyrp1b*, pigment cell progenitor marker *sox10* and cell cycle gene *pcna* as feature plots on the UMAP plot from (A). (C) Dynamic changes in subpopulations of *mitfa*^+^*aox5*^lo^ cells. Biological sample runs were downsampled to a common number of total cells so shifts in cluster proportions could be readily visualized. (D) Quantification of proportion of cells per scRNAseq sample in the melanocyte, cycling differentiation, and direct differentiation subpopulations during regeneration. P values calculated using differential proportion analysis (Farbehi et al., 2019), * p < 0.05, ** p < 0.01; *** p < 0.001; **** p < 0.0001; ns, not significant. (E) Cellular trajectories, as determined by Monocle3 (Cao et al., 2019) projected onto the *mitfa*^+^*aox5*^lo^ subcluster. Solid lines represent trajectories, with an origin in the McSC-0 subpopulation and two distinct paths through different intermediate cell subpopulations. (F) Ridge plot of the distribution of pseudotime during regeneration. Height of ridge corresponds to number of cells at that pseudotime.

These differences in gene expression between subpopulations, and dynamic subpopulation sizes, suggested that cells transit between subpopulations during regeneration. However, our analyses thus far were limited to our real-time sampling and could miss the fluidity and heterogeneity of biological process time. To further interrogate the transcriptional changes present during differentiation from McSCs to melanocytes, we utilized a latent variable, pseudotime analysis (Trapnell et al., 2014; Wolf et al., 2018). We applied the R package monocle3 to learn graph trajectories for our *mitfa^+^aox5^lo^* subgroup of cells, which includes McSC and melanocyte subpopulations (Cao *et al*., 2019). We set the McSC-0 subpopulation as pseudotime 0, an approach validated by known neural crest and melanocyte synthesis markers as well as RNA splicing mechanics as determined by RNA velocity analyses (Supplemental Figure 2E). Pseudotime analysis projected onto the *mitfa^+^aox5^lo^* subgroup of cells suggested two trajectories: one from the McSC-0 subpopulation through the direct differentiation subpopulation into melanocytes and another branching through the McSC-1 and cycling differentiation subpopulations into melanocytes (Figure 2E). Using this graph projection, we ordered the cells by biological pseudotime. McSCs are low pseudotime cells (enriched for *sox4a*, *sox10*, and no melanin synthesis genes) while melanocytes are high pseudotime cells (enriched for *tyrp1b* and *pmela*). This pseudotime dynamic mirrors what is seen *in vivo* when unpigmented *mitfa*+ cells differentiate. Visualization of pseudotime distribution differences between real-time samples reinforces the dynamic changes of McSCs during regeneration (Figure 2F). High pseudotime melanocytes were lost following ablation between day 0 and day 1, resulting in a distribution of predominantly low pseudotime McSCs at day 1. Then, as regeneration proceeded the intermediate cell states became enriched and can be visualized as medium pseudotime cells in days 2-5. Together these analyses provide a comprehensive picture of changes in McSCs transcriptomes during melanocyte regeneration which mirror real-time regeneration observations.

To assess if our approach could inform broader mechanisms governing melanocyte regeneration, we compared our zebrafish McSC signature with a mammalian McSC signature. A recent single-cell transcriptomic analysis of murine McSCs during hair follicle homeostatic cycling provided an opportunity to compare zebrafish and mammalian signatures (Infarinato *et al*., 2020). Clustering of 626 wild-type murine McSCs and melanocytes revealed four populations of previously described cells: quiescent McSCs (qMCsCs), activated McSCs (aMcSCs), cycling McSCs, and melanocytes (Supplemental Figure 3A) (Infarinato *et al*., 2020). Visualization of this structure reveals similarities to the previously computed zebrafish clustering, where non-cycling McSCs are separated from mature melanocytes by intermediate cell populations. To assess similarities in gene expression we calculated marker genes for each zebrafish and murine subpopulation, and then visualized conserved gene signatures via heatmaps (Supplemental Figure 3B). As murine McSCs progress from qMcSCs to differentiated melanocytes they lose expression of *Zeb2*, *Pax3*, and AP-1 FOS and JUN family subunits, while upregulating melanin synthesis genes such as *Tyrp1* and *Oca2*. These changes mirror the transcriptional profiles found in our zebrafish dataset. Lastly, to visualize correlation between zebrafish and murine populations we calculated cluster-specific differentially expressed genes, mapped these to murine orthologs, and then scored the murine populations with these zebrafish signatures (Supplemental Figure 3C). Through these comparisons, we find that our McSC-0 and McSC-1 subpopulations are most similar to qMcSCs. The zebrafish direct differentiation subpopulation, much like when it is compared to other zebrafish subpopulations, shares aspects of murine McSC and more differentiated subpopulations. The zebrafish cycling differentiation subpopulation most strongly overlaps with murine cycling McSC subpopulation. Reassuringly, mature pigment producing melanocytes in each dataset are most like each other. These signature scores reflect the positional similarities we see in UMAP visualization where qMcSCs and McSC-0 and McSC-1 are most stem-like and least differentiated. Overall, these signatures support a shared gene signature of differentiation during McSC activation, supporting the use of zebrafish to uncover conserved signaling pathways governing melanocyte regeneration.

### Signaling pathway dynamics in McSCs uncover the KIT signaling axis as a candidate regulator of melanocyte regeneration

To identify signaling pathways governing melanocyte regeneration we first analyzed potential receptor-ligand interactions using NicheNetR (Browaeys et al., 2020). In brief, NicheNetR calculates transcriptional changes in a target “receiver” cell population between time samples. It uses these differentially expressed genes to predict ligands responsible for the observed behavior. Finally, it matches the expression of these ligands on potential “sender” cells with cognate receptors on the receiver cell subpopulation. We designated our identified McSC subpopulations as receiver cells and all other cell types as potential sender cells. We compared gene expression in our regenerating McSCs on days 2, 3 and 5 to McSC gene expression in the unactivated day 0 controls. We then filtered down top scoring ligand/receptor pairs to NicheNetR’s ‘bona fide’ literature-supported pairs. This approach identified several signaling systems with known roles in melanocyte biology, including KITLG/KIT, ASIP/MC1R, EDN3/EDNRB and NRG1/ERBB2 (Figure 3A) (Chou *et al*., 2013; Hultman *et al*., 2009; Li et al., 2017; Yamada et al., 2013). These signaling systems function during melanocyte development, and our data indicate they are also reactivated during melanocyte regeneration. To more deeply understand how such reactivation would modulate regeneration, we focused on KITLG/KIT signaling. In zebrafish the KIT receptor ortholog, *kita*, is necessary for the survival and migration of melanocytes as they emerge from the neural crest (Rawls and Johnson, 2003) and has been implicated in establishment of larval McSCs (O’Reilly-Pol and Johnson, 2013; Yang et al., 2004), but whether the KITLG/KIT pathway is reactivated during regeneration and how it governs McSC fates have not been addressed.

**Figure 3.**
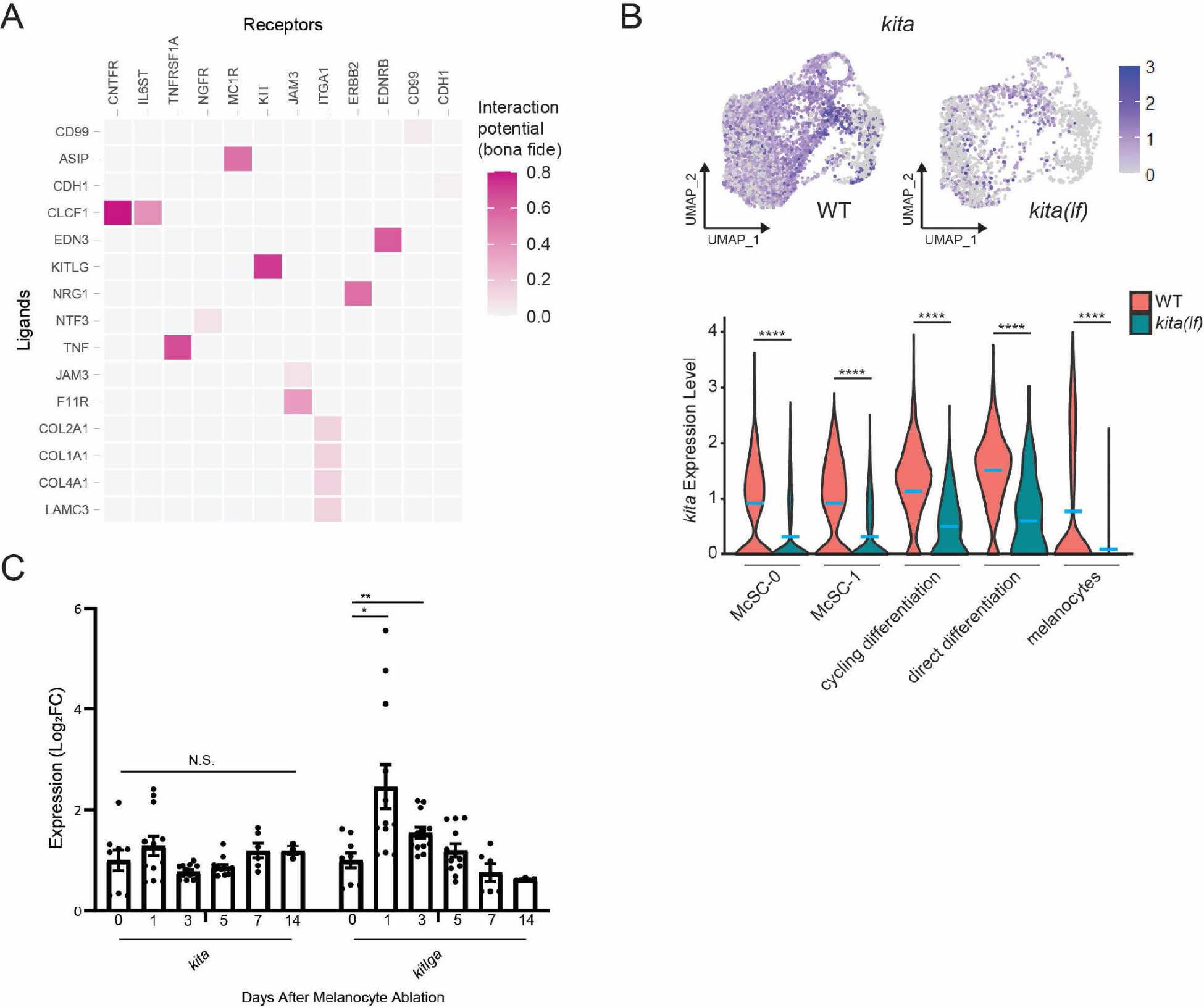
NicheNet and transcriptional analyses implicate the KIT signaling axis as a dynamic regulator of melanocyte regeneration. (A) Heatmap of NicheNetR-identified ligand/receptor pairs indicating interaction potential between McSC receiver cells and other sender cells sampled by scRNAseq. Ligand/receptor pairs were restricted to literature-supported pairs (Browaeys *et al*., 2020). (B) Top, feature and, bottom, violin plots of *kita* expression in *mitfa*^+^*aox5*^lo^ cells from wild-type and *kita(lf)* strains. Mean gene expression represented by cyan bars. P values calculated by Wilcoxon rank sum test, **** p < 0.0001. (C) qRT-PCR of *kita* and *kitlga* expression in zebrafish skin following melanocyte destruction. Three biological replicates were performed for each time point. Data are shown as mean + SEM. P values calculated by Student’s t-test, * p < 0.05 and ** p < 0.01; ns, not significant.

We analyzed expression of *kita* and its ligand *kitlga* during regeneration. *kita* receptor was expressed in McSC, cycling differentiation and direct differentiation subpopulations (Figure 3B). To test for dynamism in the KIT signaling axis, we sought to assay *kita* receptor and *kitlga* ligand expression following melanocyte destruction (Figure 3C). From bulk skin samples, levels of *kita* receptor did not change dramatically during regeneration. By contrast, *kitlga* ligand levels changed such that there was an increase of *kitlga* expression shortly following melanocyte destruction, which then tapered downwards as regeneration proceeded. We analyzed scRNAseq data to determine cell types that express *kitlga*. Thanks to trace numbers of *mitfa*-negative cells that were analyzed as part of our scRNAseq, we were able to observe *kitlga* expression in fibroblasts and keratinocytes, mirroring scRNAseq expression patterns from human cells (Supplemental Figure 4A,B) (Joost et al., 2020). Previous studies have shown that KIT has a critical role in mediating lineage decisions in melanocyte development as well as melanocyte homeostasis in the murine hair follicle (Botchkareva et al., 2003; Botchkareva et al., 2001; Geissler et al., 1988; Rawls and Johnson, 2003). KITLG/KIT signaling also has known roles in human melanoma as well as murine and zebrafish melanoma models (Beadling et al., 2008; Santoriello et al., 2010). Our data indicate that KITLG/KIT signaling is reactivated in response to injury, making this signaling axis a leading candidate as a regulator of McSC fates during regeneration.

### kita receptor and kit ligand loss-of-function mutants impair direct differentiation of McSCs during melanocyte regeneration

To directly test the role of KITLG/KIT signaling predicted by transcriptomics we measured melanocyte regeneration in *kita(lf)* and *kitlga(lf)* animals. Whereas wild-type animals regenerated their full contingent of stripe melanocytes within two weeks of neocuproine-induced melanocyte destruction, *kita(lf)* and *kitlga(lf)* mutants regenerated 51.4% and 65.2% of their melanocytes, respectively, indicating a requirement for KIT signaling during regeneration (Figure 4A,B). Previous work established that zebrafish regeneration melanocytes arise from one of two sources: direct differentiation of McSCs or, less frequently, asymmetric divisions of McSCs in which one of two daughter cells undergoes differentiation (Iyengar *et al*., 2015). To determine whether one or both McSC fates were defective in *kita* mutants we sequenced 24,724 transcriptomes from *mitfa:nlsEGFP*-sorted cells from regenerating *kita(lf)* mutants. Integrated clustering of cells from *kita(lf)* mutant and wild-type strains revealed conservation of the cell types found in both strains (Supplemental Figure 5A). Additionally, analysis of *kita* expression showed that *kita(lf)* mutants did indeed express lower levels of *kita* transcripts, likely due to nonsense-mediated decay tied to a premature stop codon encountered after the frameshift mutation in the *kita(b5)* allele used in our study (Figure 3B) (Parichy et al., 1999a). We analyzed scRNAseq data and performed *in vivo* lineage tracing to determine the basis for the regeneration defect in *kita/kitlga* animals. Trajectory analysis using monocle3 revealed that the pathway from McSC subpopulations through the direct differentiation subpopulation to melanocytes was absent in *kita(lf)* mutants (Figure 4C). This absence stemmed from the lack of an increase in the direct differentiation subpopulation during regeneration (Figure 4D). The cycling differentiation subpopulation was also affected, with a smaller increase during regeneration observed as compared to the wild type. To assess if these differences resulted from lineage defects during regeneration, we performed lineage tracing in *kita(lf)* mutants containing the *mitfa(nls:EGFP)* transgene. Lineage tracing revealed a greater than 8-fold decrease in McSCs undergoing direct differentiation (Figure 4E). Asymmetric divisions were also decreased by 2-fold. Lineage tracing of *kitlga(lf); Tg(mitfa:nlsEGFP)* mutants showed similar defects as those observed in *kita(lf)* mutants (Figure 4E). Together, these data elucidate paths and signaling requirements for the two fates adopted by McSCs to regenerate new melanocytes. In one fate McSCs directly differentiate through an intermediate cell state before becoming melanocytes. This fate has a strong requirement for KIT signaling. In the other fate, McSCs take part in asymmetric divisions to generate new melanocytes. This fate path is characterized by co-expression of differentiation and cell cycle genes and is also affected, albeit less so, by defects in KIT signaling.

**Figure 4.**
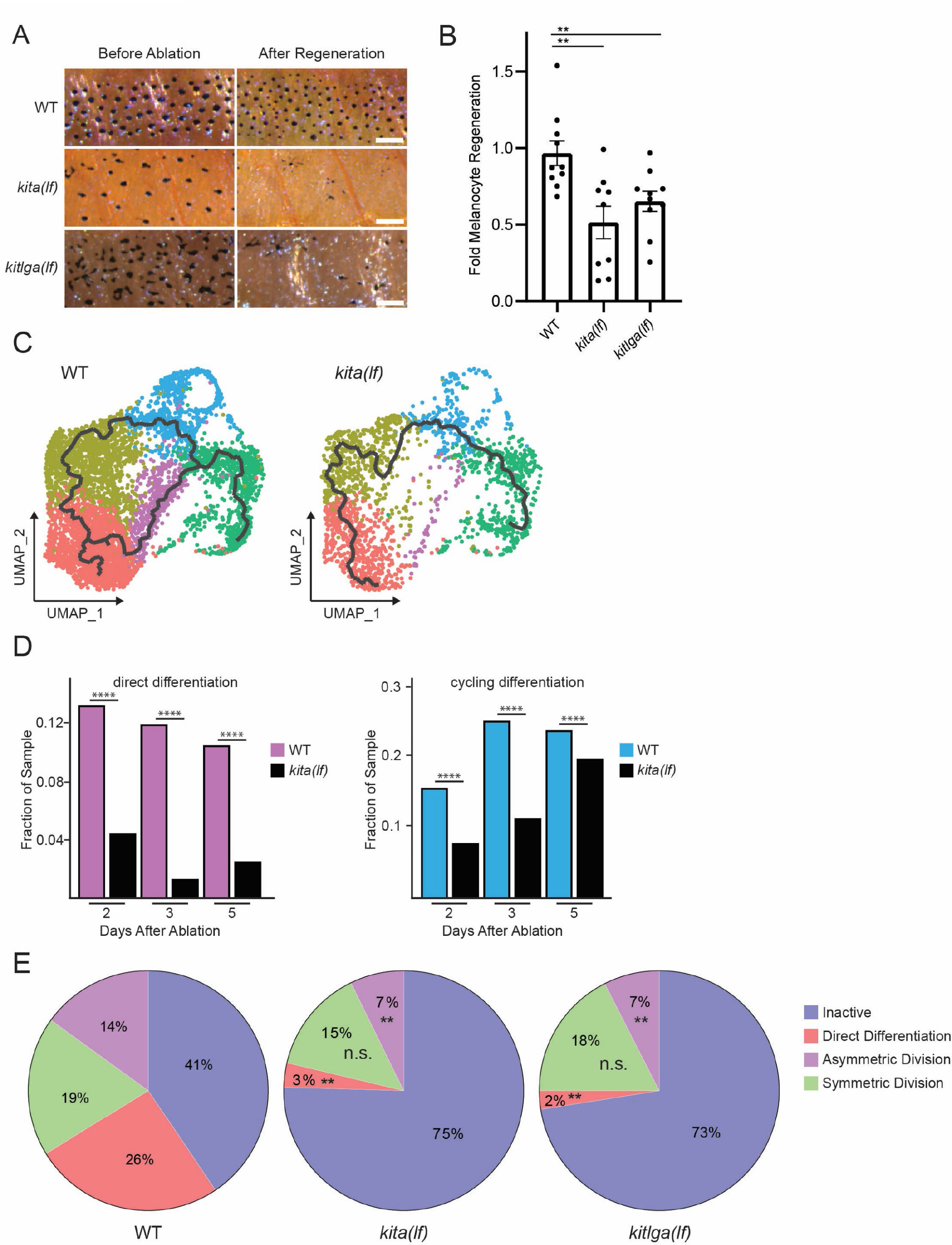
*kita* receptor and *kitlga* ligand loss-of-function mutants have impaired melanocyte regeneration. (A) Brightfield images of the melanocyte stripe before melanocyte ablation and after melanocyte regeneration in wild-type, *kita(lf)*, and *kitlga(lf)* zebrafish strains. Scale bar =200µm. (B) Quantification of melanocyte regeneration in wild-type, *kita(lf)* and *kitlga(lf)* strains. Mean + SEM is shown; WT n = 10, *kita(lf)* = 9, *kitlga(lf)* = 11 fish. P values calculated by Student’s t-test, ** p < 0.01. (C) Cellular trajectories, using Monocle3, of *mitfa*^+^*aox5*^lo^ cells from wild-type zebrafish (left) and *kita(lf)* mutants (right). Solid lines represent trajectories, with an origin in the McSC-0 subpopulation and terminus in the melanocyte subpopulation. (D) Quantification of McSCs and melanocyte lineage subpopulations in regeneration samples (days 2, 3, 5 post-ablation). Reductions in *kita(lf)* zebrafish were observed in direct differentiation (3.8-fold) and cycling differentiation (2.1-fold) cell subpopulations. Increases in McSC-0 (1.1-fold) and McSC-1 (1.7-fold) cell subpopulations were also observed. (D) Comparison of proportion of WT and *kita(lf)* cells per scRNAseq sample in the cycling differentiation and direct differentiation subpopulations during regeneration reveals fewer *kita(lf)* cells going through differentiation. P values calculated using differential proportion analysis (Farbehi *et al*., 2019), **** p < 0.0001; ns, not significant. (E) Fates of McSCs following single-cell lineage tracing of wild-type, *kita(lf)* and *kitlga(lf)* strains. Mean percentage of traced McSCs in a fate are shown; wild type n = 4, *kita(lf)* = 4, *kitlga(lf)* = 4. P values calculated by Student’s t-test, ** p < 0.01, ns, not significant.

### Kit-mediated differentiation depends on MAPK pathway activity

The defect in regeneration caused by loss of *kita* receptor or *kitlga* ligand suggests that signaling downstream of the KIT receptor is required for proper McSC differentiation during regeneration. One well-documented downstream pathway is the MAPK pathway, with ERK being a known controller of the melanocyte lineage master regulator *mitfa* (Hemesath et al., 1998; Levy et al., 2006; Wellbrock and Arozarena, 2015; Wellbrock et al., 2008). Accordingly, we hypothesized that if KITLG/KIT signaling is important in regeneration, then ERK activity would be upregulated in McSCs following melanocyte destruction. To test this hypothesis, we utilized *in vivo* imaging with a kinase translocation reporter, ERKKTR-mClover (Mayr et al., 2018; Regot et al., 2014). In this system ERK activation via phosphorylation drives the ERKKTR-mClover fusion protein from a nuclear to cytosolic localization. These shifts in subcellular localization allow quantification of ERK activity and, by extension, MAPK pathway activity. We generated a MAPK activity reporter that drives ERKKTR-mClover expression from a melanocyte lineage-specific *mitfa* promoter and injected this reporter construct into a *Tg(mitfa:nlsmCherry)* strain to enable accurate measurement of nuclear/cytosolic intensity. We first observed that McSCs showed lower ERK activity than mature melanocytes (Figure 5A,B), indicating that ERK activity in McSCs is relatively low under homeostatic conditions.

**Figure 5.**
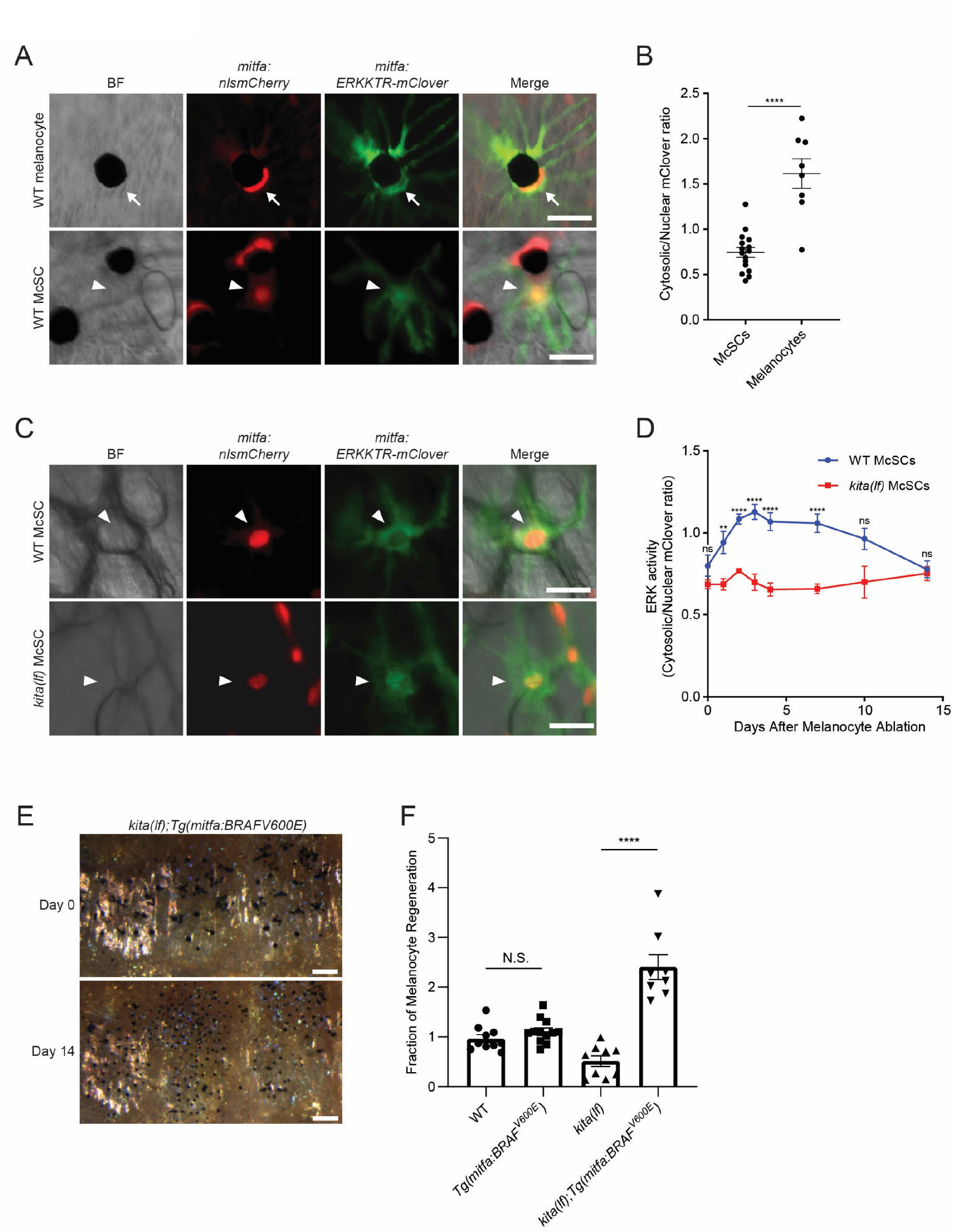
*kitlga/kita* signaling during melanocyte regeneration acts through the MAPK pathway. (A) Images of ERKKTR-mClover localization in a representative mature melanocyte (top, arrow) and McSC (bottom, arrowhead) in uninjured wild-type animals. Scale bar = 30 µm. (B) Quantification of ERK activity in McSCs and melanocytes based on ERKKTR-mClover localization. Mean ± SEM is shown; McSCs n = 16, melanocytes n = 8. (C) Images 3 days post-ablation of ERKKTR-mClover location in representative McSCs (arrowheads) in wild-type (top) and *kita(lf)* (bottom) animals. Scale bar = 30µm. (D) Quantification of ERK activity in McSCs prior to and during melanocyte regeneration. For each data point the average cytosolic/nuclear ratio of at least 6 cells ± SEM is shown. (E) Brightfield images of the melanocyte stripe before (top) and after (bottom) regeneration in *kita(lf); Tg(mitfa:BRAFV600E)* mutants. Scale bar = 200µm. (F) Quantification of melanocyte regeneration in *kita(lf);Tg(mitfa:BRAFV600E)* and control animals. Mean + SEM is shown; wild type n = 10, *Tg(mitfa:BRAFV600E)* = 12, *kita(lf)* = 9, *kita(lf);Tg(mitfa:BRAFV600E)* = 8 fish. P values calculated by Student’s t-test, ** p < 0.01, **** p < 0.0001; ns, not significant.

To further understand the role of MAPK signaling in regeneration we assayed ERKKTR-mClover localization following melanocyte ablation. Following ablation, ERKKTR-mClover signal in wild-type McSCs shifted to an increased cytosolic localization, indicative of an increase in MAPK activity during regeneration (Figure 5C,D). Next, to test whether this increase was dependent on KITLG/KIT signaling, we injected the *mitfa:ERKKTR-mClover* reporter into a *kita(lf); Tg(mitfa:nlsmCherry)* strain. McSCs in *kita(lf)* mutants displayed similar levels of ERK activity in homeostatic conditions as compared to McSCs in wild-type animals (Figure 5C,D). However, following melanocyte injury the *kita(lf)* animals showed no cytosolic translocation of ERKKTR-mClover, indicating a failure to upregulate MAPK activity in the absence of KIT signaling (Figure 5C,D). These results indicate that McSCs upregulate MAPK activity during regeneration, and blockade of KIT signaling impedes this upregulation.

If KIT-mediated MAPK signaling is required for direct differentiation during regeneration, then rescue of downstream MAPK signaling in *kita(lf)* mutants should result in rescue of the regeneration phenotype. To test this hypothesis, we utilized a constitutively active RAF, BRAF^V600E^, a component of the MAPK pathway downstream of KIT but upstream of ERK (Davies et al., 2002; Patton et al., 2005). Introduction of the overactive BRAF had little effect in wild-type animals, but rescued regeneration in *kita(lf)* animals resulting in a melanocyte density that is comparable to that of wild-type animals (Figure 5E,F; Supplemental Figure 5B). These results show that functional KIT signaling through the MAPK pathway is necessary and sufficient for proper melanocyte regeneration.

### scRNAseq identifies *mitfa^+^aox5^hi^* cells that undergo divisions following melanocyte destruction

Multiple rounds of melanocyte injury can be performed in zebrafish without diminishing the capacity to regenerate. This is due to a subset of McSCs that divide symmetrically to maintain the pool of McSCs. Previous studies found that daughters of these symmetric divisions that arise after one round of injury are capable of differentiation following subsequent injury, indicating a link between self-renewal and eventual differentiation (Iyengar *et al*., 2015). Given their central role in melanocyte regeneration, we sought to understand characteristics of McSCs that undergo symmetric divisions through our scRNAseq dataset.

To begin, we investigated our scRNAseq dataset for cells that entered the cell cycle in response to melanocyte injury. As previously mentioned, a subpopulation of *mitfa^+^aox5^lo^* cells enter the cell cycle and express differentiation markers. Trajectory analysis indicates some of these cells ultimately differentiate into melanocytes, suggesting that the dividing cells may be ones that undergo asymmetric divisions and generate a differentiated daughter cell during regeneration. In addition to these cells, we identified separate *mitfa^+^aox5^hi^* cycling cells that clustered into two subpopulations, one *pcna*-enriched and high in S phase score (*mitfa^+^aox5^hi^* S phase) and the other *cdk1*-enriched and high in G2/M score (*mitfa^+^aox5^hi^* G2/M phase) (Figure 6A,B, Supplemental Figure 6A). Neither subpopulation expressed the melanization genes *tryp1b* and *pmela* (Supplemental Figure 6B). Plotting these two subpopulations from each sampled time point revealed very few cells prior to melanocyte injury (Day 0), a robust increase during regeneration (Days 1,2,3,5) and a diminution when regeneration was mostly complete (Day 10) (Figure 6C,D). The cell cycle phase scoring indicated a clockwise cellular trajectory from *mitfa^+^aox5^hi^* S phase to *mitfa^+^aox5^hi^* G2/M phase subpopulations, and RNA velocity analysis suggested that an adjacent subpopulation (*mitfa^+^aox5^hi^* pre-cycle) was an input to *mitfa^+^aox5^hi^* S phase and a separate adjacent subpopulation (*mitfa^+^aox5^hi^* post-cycle) was an output from *mitfa^+^aox5^hi^* G2/M phase (Figure 6E). As was evident from their separate clustering, *mitfa^+^aox5^hi^* pre-cycle and *mitfa^+^aox5^hi^* post-cycle subpopulations had differential expression of several genes, suggesting that the cell divisions involved are not limited to self-renewal. Interestingly, as compared to other subpopulations, genes differentially upregulated in the *mitfa^+^aox5^hi^* post-cycle subpopulation were associated with McSC-0 and McSC-1 subpopulations, suggesting a possible relationship between these subpopulations (Figure 6F).

**Figure 6.**
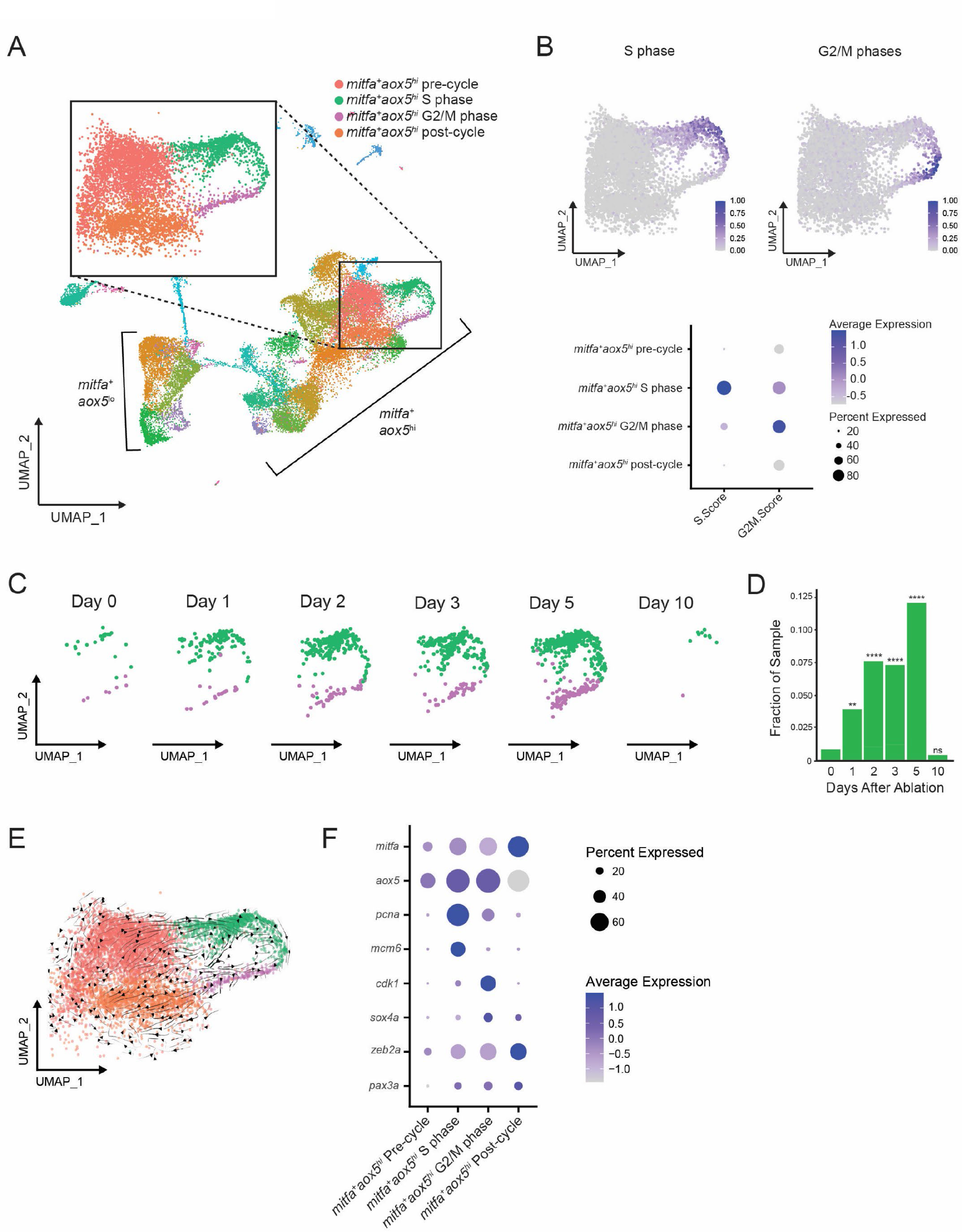
A subpopulation of *mitfa*^+^*aox5*^hi^ cells divides and expands during regeneration. (A) UMAP of all cells sampled from wild-type animals with enlargement of cycling and adjacent subpopulations found in the large group of *mitfa*^+^*aox5*^hi^ cells. (B) Top, feature and, bottom, dot plot of S phase and G2/M phase cell cycle scores of cells highlighted in (A). Cell cycle scores were calculated using Seurat’s ‘CellCycleScoring’ module and zebrafish orthologs of the cell-cycle genes outlined by Tirosh et al., 2016 (C) UMAPs of cycling cells prior to and during regeneration. Biological sample runs were down-sampled to a common number of total cells so shifts in clusters could be readily visualized. (D) Quantification of proportion of cells per scRNAseq sample in the *mitfa*^+^*aox5*^hi^ cycling subpopulations (*mitfa^+^aox5^hi^* S phase and *mitfa^+^aox5^hi^* G2/M subpopulations combined) during regeneration. P values calculated using differential proportion analysis (Farbehi *et al*., 2019), ** p < 0.01; **** p < 0.0001; ns, not significant. (E) RNmarker and coloring represents averageA splicing mechanics of pre-cycle, cycling, and post-cycle *mitfa*^+^*aox5*^hi^ subpopulations scored using velocity embedding (velocyto; (La Manno et al., 2018)) and visualized through velocity stream analysis (scvelo; (Bergen et al., 2020)) on the UMAP plot inset from Figure 6A. (F) Dot plot of pigment cell and cell cycle marker genes differentially expressed across pre-cycle, cycling, and post-cycle *mitfa*^+^*aox5*^hi^ subpopulations during melanocyte regeneration. Dot sizes represent percentage of cells in the cluster expressing the marker and coloring represents average expression.

The dynamics of cell division and absence of melanocyte differentiation genes make this subpopulation a candidate for McSCs that undergo symmetric divisions during regeneration. However, these cells express higher levels of *aox5*, which is a gene that is also found in differentiated xanthophores. We considered the possibility that these *mitfa^+^aox5^hi^* cycling cells were xanthophore lineage cells cycling in response to ablation of mature xanthophores by neocuproine. To investigate this possibility, we traced inter-stripe xanthophores prior to and following neocuproine treatment. Consistent with previous reports (O’Reilly-Pol and Johnson, 2008) we observed no xanthophore ablation caused by neocuproine, and out of 65 inter-stripe xanthophores traced following neocuproine ablation we observed zero cell divisions (Supplemental Figure 7). In the absence of xanthophore ablation and regeneration, we directly tested the hypothesis that the symmetrically dividing McSCs in the melanocyte stripe were *mitfa^+^aox5^hi^* cells evident in scRNAseq profiling. We combined our *mitfa:nlsmCherry* reporter with an *aox5:PALM-EGFP* reporter (Eom et al., 2015) and investigated dual positive cells in the melanocyte stripe. All of the unpigmented *mitfa*-positive cells in the melanocyte stripe also expressed *aox5:PALM-EGFP*. The dual positive cells exhibited heterogeneity in their *aox5:PALM-EGFP* expression consistent with one subpopulation being *aox5^hi^* and another subpopulation *aox5^lo^* (Figure 7A,B). To determine if *mitfa^+^aox5^hi^* cells correspond to McSCs that undergo symmetric divisions during regeneration, we tracked dual positive cells following melanocyte ablation (Figure 7C). Cells that underwent symmetric divisions during regeneration exhibited high *aox5^+^* intensities before melanocyte regeneration (Figure 7D). Taken together, these results suggest that the cycling *mitfa^+^aox5^hi^* cells from our scRNAseq correspond to McSCs in the melanocyte stripe that undergo symmetric divisions following melanocyte destruction.

**Figure 7.**
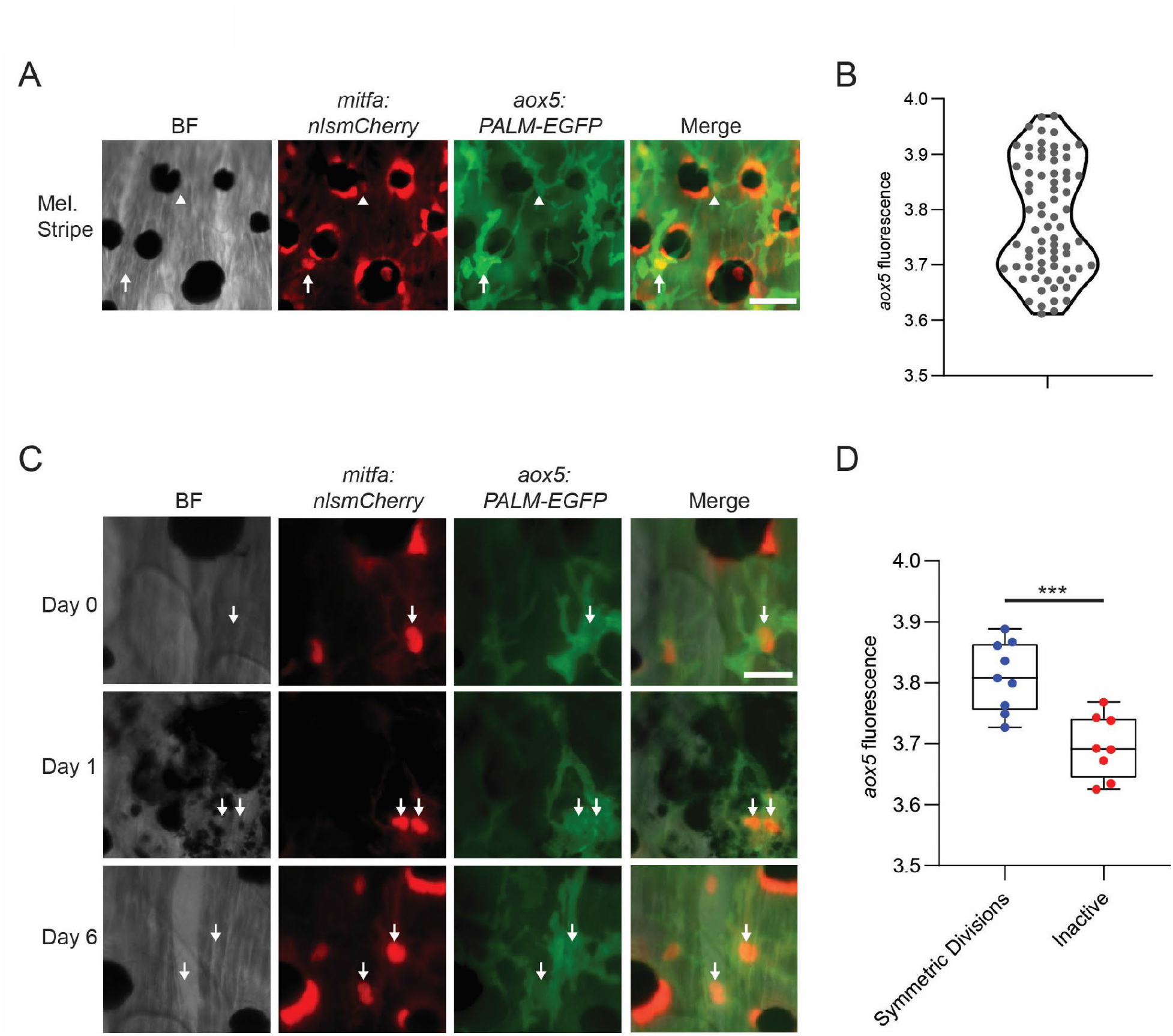
*aox5* expression predicts *in vivo* McSC cell fate. (A) Representative image of McSCs expressing different levels of *aox5* promoter-driven PALM-EGFP. *mitfa^+^aox5^hi^* (arrowhead), *mitfa^+^aox5^lo^* (arrow). Scale bar = 50µm. (B) Quantification of PALM-EGFP fluorescence intensity indicates groups of McSCs that express lower and higher levels of *aox5*. Intensity values are log normalized. (C) Images of an *mitfa*^+^*aox5*^hi^ cell that underwent mitosis following melanocyte destruction. Scale bar = 30µm. (D) Comparison of *aox5* intensity in cells that underwent symmetric divisions or remained inactive. P values calculated by Student’s t-test, *** p < 0.001.

## Discussion

Stem cells that are activated in tissue regeneration and homeostasis have the capacity to generate newly differentiated cells and maintain a pool of stem cells. In this study we combined scRNAseq with single-cell lineage tracing and other analyses to determine how this occurs during regeneration of melanocyte stripes in zebrafish. Interestingly, we found that McSC heterogeneity and fate-specific signaling systems underlie a coordinated regeneration process.

McSC heterogeneity is illustrated, in part, by the two subpopulations of *mitfa^+^aox5^lo^* McSCs, McSC-0 and McSC-1. McSC-0 and McSC-1 are transcriptionally distinct and are present in unperturbed animals. RNA velocity and cell trajectory analyses indicates that these subpopulations are activated upon melanocyte ablation and that each supplies different intermediate subpopulations during the regenerative process. McSC-1 cells feed primarily into the cycling differentiation subpopulation, whereas McSC-0 cells feed into the direct differentiation subpopulation. Additionally, these analyses suggest that McSC-0 is a more truncal subpopulation that can also backfill the McSC-1 subpopulation during regeneration. The benefit to having distinct subpopulations of McSCs is unclear, although it appears to enable regeneration of new melanocytes by two different routes, the differential regulation of which may be important for regeneration to occur proficiently under different circumstances. Regardless, the resolution afforded by scRNAseq indicates that the McSC-0 and McSC-1 subpopulations are present in unperturbed animals and primed to adopt different fates when activated in response to melanocyte injury.

Heterogeneity is also evident by the additional *mitfa^+^aox5^hi^* subpopulation that enters the cell cycle following melanocyte ablation. There are several reasons to think that this is an McSC subpopulation. Firstly, these cells were activated in response to specific ablation of melanocytes. Secondly, direct observations and lineage tracing showed that a subset of *mitfa^+^aox5^hi^* cells divided symmetrically during melanocyte regeneration, and we showed previously that such symmetrically dividing cells could generate new melanocytes in subsequent cycles of melanocyte regeneration (Iyengar *et al*., 2015). Thirdly, although this subpopulation clusters more closely to xanthophores, we showed that xanthophores are not ablated by the melanocyte-ablating drug neocuproine and this *mitfa^+^aox5^hi^* subpopulation does not make new xanthophores following neocuproine treatment. Fourthly, the properties of this subpopulation are reminiscent of multipotent pigment progenitor cells during development, in that they express high levels markers that are expressed by different types of terminally-differentiated cells (Budi et al., 2008; Dooley et al., 2013; Singh et al., 2016). These reasons support a conclusion that the *mitfa^+^aox5^hi^* cells that enter the cell cycle during melanocyte regeneration correspond to cells that are observed *in vivo* to symmetrically divide during melanocyte regeneration. Whether all of the cells that undergo symmetric divisions come from a pool of *mitfa^+^aox5^hi^* cells is unclear; we did not observe *mitfa^+^aox5^lo^* cells undergo symmetric divisions *in vivo* but we cannot exclude the possibility that some small portion of *mitfa^+^aox5^lo^* cycling differentiation subpopulation also divides symmetrically during regeneration.

Interesting questions arise pertaining to the potential relationship between a subpopulation of dividing *mitfa^+^aox5^hi^* cells and the McSC-0 and McSC-1 subpopulations. Previous fate mapping and lineage tracing indicated that McSCs that divided symmetrically maintained the pool of McSCs for subsequent regeneration cycles. Consequently, if the McSCs we observed to symmetrically divide *in vivo* were derived from *mitfa^+^aox5^hi^* mother cells, then do some of these daughter cells ultimately transition to *mitfa^+^aox5^lo^* McSC-0 or McSC-1 cells? That would be the expectation based on our current understanding of stripe melanocyte regeneration, and it would be similar to what occurs in some tissues, such as the lung, where multipotent epithelial stem cells symmetrically divide before giving rise to more lineage-committed stem cells (McQualter *et al*., 2010). Supporting this notion is the presence of the *mitfa^+^aox5^hi^* post-cycle cluster, in which the stem cell marker *sox4a*, the neural crest genes *zeb2a* and *pax3a* as well as other genes characteristic of McSC-0 and McSC-1 supopulations are upregulated. However, our current results do not indicate when a transition from *mitfa^+^aox5^hi^* post-cycle cells to McSC-0 or McSC-1 cells would occur and how long it would take for cells to transition. It is possible that this transition is slow or occurs at time points not sampled in the current study. Another question is whether the symmetric divisions of McSCs we observed in vivo are truly symmetric. We have described them as such because both daughter cells remain undifferentiated in vivo. However, if these cells are the same as the dividing *mitfa^+^aox5^hi^* cells from scRNAseq, we cannot distinguish between the possibilities that: a) symmetric divisions occur to yield two *mitfa^+^aox5^hi^* post-cycle daughters, or b) asymmetric divisions occur to yield an *mitfa^+^aox5^hi^* post-cycle daughter and an *mitfa^+^aox5^hi^* pre-cycle daughter. The second possibility includes some degree of self-renewal, which is likely to occur given the apparent unlimited nature of zebrafish melanocyte regeneration. One last question is whether a transition to McSC-0 or McSC-1 cells is the only fate available to *mitfa^+^aox5^hi^* dividing cells. As noted above, the *mitfa^+^aox5^hi^* daughter cells from symmetric divisions are similar to multipotent pigment progenitor cells in development. It is possible that they are capable of ultimately adopting a melanocyte fate or that or the other pigment cell types, xanthophores or iridophores. These three questions are beyond the scope of the current study but nonetheless are worth considering as they help to understand the complexity of pigment cell regeneration in zebrafish and stem-cell-mediated regeneration in general.

Our investigation of KIT signaling sheds light on how the process of melanocyte regeneration is controlled. KIT signaling has well-known roles in melanocyte development in promoting the differentiation, survival and migration of melanocytes. Here, we have found that KIT signaling is reactivated during melanocyte regeneration to enable McSCs to generate new melanocytes. Specifically, shortly after melanocyte ablation we observed increased expression of the *kitlga* ligand, and the increased expression of *kitlga* was followed shortly thereafter by an increase in MAPK signaling in McSCs. KIT receptor mutants showed a profound reduction in cells in the direct differentiation subpopulation and a less marked, but nonetheless significant, reduction in the cycling differentiation subpopulation. In these mutants there were no changes in the *mitfa^+^aox5^hi^* cycling subpopulation. Interestingly, MAPK signaling was decreased in all McSCs we measured in KIT receptor mutants. This suggests that, while all subpopulations of McSCs might upregulate KIT-dependent MAPK signaling during regeneration, these subpopulations are differentially dependent on this activity for executing their fates during regeneration, i.e. McSC-0 cells fated to undergo direct differentiation are most dependent, whereas *mitfa^+^aox5^hi^* cells fated to undergo symmetric divisions are least dependent.

Overall, our study has found unexpected heterogeneity of McSC subpopulations, defined new intermediate subpopulations involved in regeneration, and discovered cellular trajectories that link these subpopulations during the regeneration process. Along with providing a greater understanding of melanocyte regeneration in zebrafish and tissue regeneration in general, these findings are relevant to pigmentary disorders and melanoma. Treatments for vitiligo, an immune-mediated destruction of melanocytes, and other pigmentary disorders stimulate McSC activity, and our results highlight cellular transitions and signaling pathways regulating these transitions that could be manipulated to facilitate pigment recovery. In melanoma, stem cell-like identities are implicated in tumor maintenance and drug resistance. A more detailed understanding of McSCs and the pathways that govern their differentiation could inform fundamental studies of melanoma growth and therapy.

## Methods

Key Resources Table

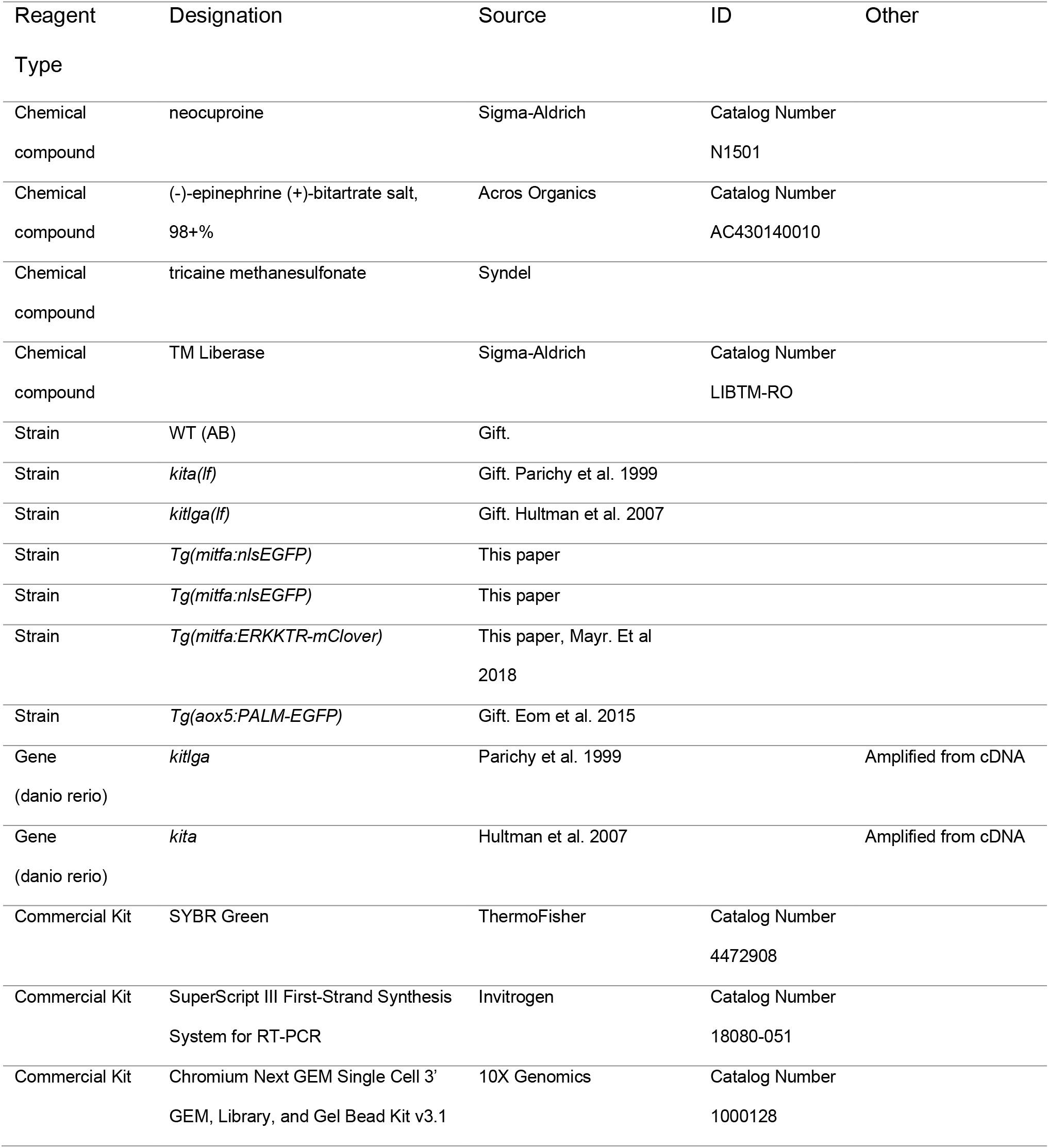
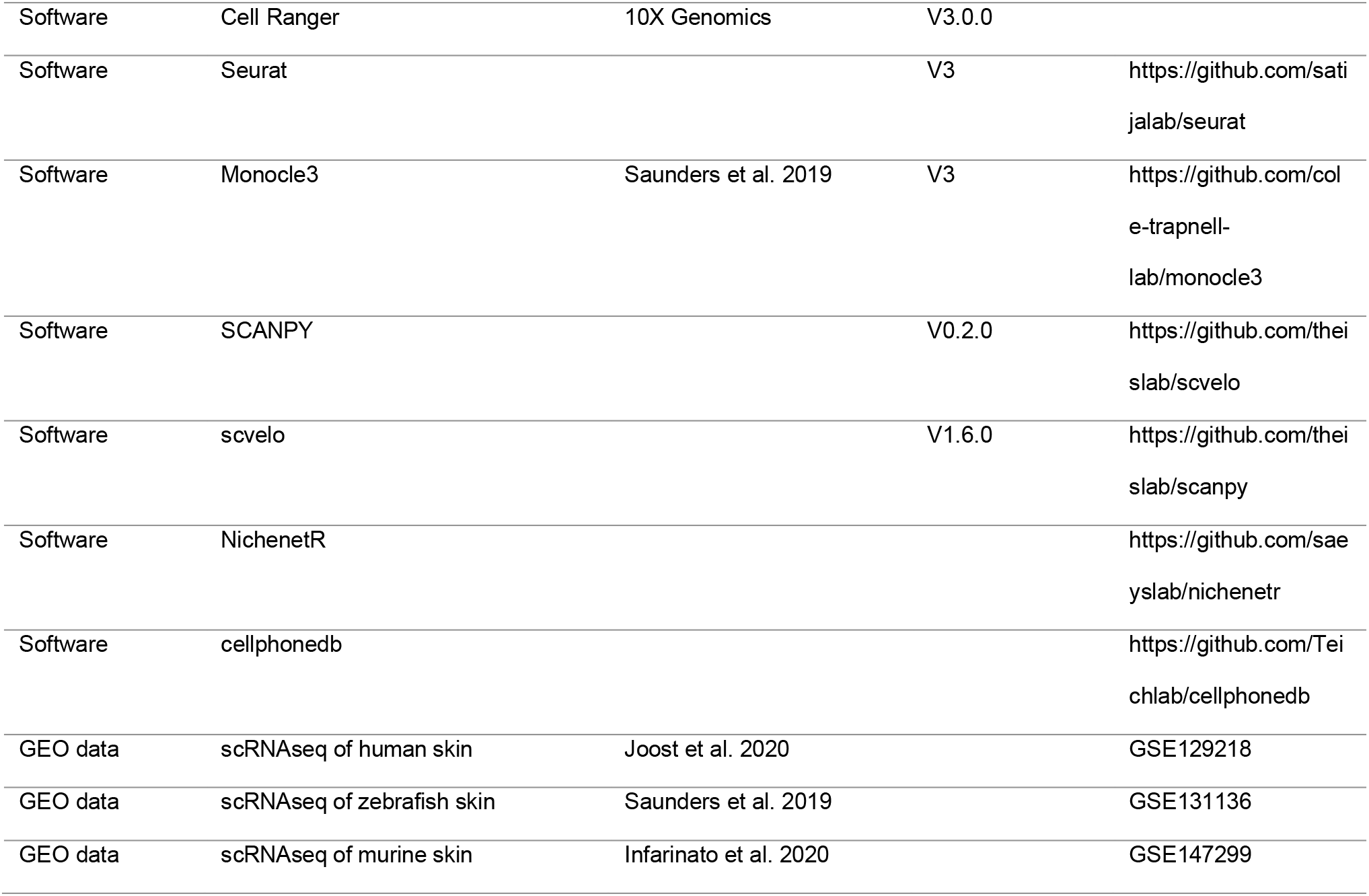

| Reagent<br>Type | Designation | Source | ID | Other |
| --- | --- | --- | --- | --- |
| Chemical<br>compound | neocuproine | Sigma-Aldrich | Catalog Number<br>N1501 |  |
| Chemical<br>compound | (-)-epinephrine (+)-bitartrate salt,<br>98+% | Acros Organics | Catalog Number<br>AC430140010 |  |
| Chemical<br>compound | tricaine methanesulfonate | Syndel |  |  |
| Chemical<br>compound | TM Liberase | Sigma-Aldrich | Catalog Number<br>LIBTM-RO |  |
| Strain | WT (AB) | Gift. |  |  |
| Strain | *kita(lf)* | Gift. Parichy et al. 1999 |  |  |
| Strain | *kitlga(lf)* | Gift. Hultman et al. 2007 |  |  |
| Strain | *Tg(mitfa:nlsEGFP)* | This paper |  |  |
| Strain | *Tg(mitfa:nlsEGFP)* | This paper |  |  |
| Strain | *Tg(mitfa:ERKKTR-mClover)* | This paper, Mayr. Et al<br>2018 |  |  |
| Strain | *Tg(aox5:PALM-EGFP)* | Gift. Eom et al. 2015 |  |  |
| Gene<br>(danio rerio) | *kitlga* | Parichy et al. 1999 |  | Amplified from cDNA |
| Gene<br>(danio rerio) | *kita* | Hultman et al. 2007 |  | Amplified from cDNA |
| Commercial Kit | SYBR Green | ThermoFisher | Catalog Number<br>4472908 |  |
| Commercial Kit | SuperScript III First-Strand Synthesis<br>System for RT-PCR | Invitrogen | Catalog Number<br>18080-051 |  |
| Commercial Kit | Chromium Next GEM Single Cell 3'<br>GEM, Library, and Gel Bead Kit v3.1 | 10X Genomics | Catalog Number<br>1000128 |  |
| Software | Cell Ranger | 10X Genomics | V3.0.0 |  |
| Software | Seurat |  | V3 | https://github.com/satijalab/seurat |
| Software | Monocle3 | Saunders et al. 2019 | V3 | https://github.com/cole-trapnell-lab/monocle3 |
| Software | SCANPY |  | V0.2.0 | https://github.com/theislab/scvelo |
| Software | scvelo |  | V1.6.0 | https://github.com/theislab/scanpy |
| Software | NichenetR |  |  | https://github.com/saeyslab/nichenetr |
| Software | cellphonedb |  |  | https://github.com/Teichlab/cellphonedb |
| GEO data | scRNAseq of human skin | Joost et al. 2020 |  | GSE129218 |
| GEO data | scRNAseq of zebrafish skin | Saunders et al. 2019 |  | GSE131136 |
| GEO data | scRNAseq of murine skin | Infarinato et al. 2020 |  | GSE147299 |

### Fish stocks and husbandry

Fish stocks were maintained at 28.5°C on a 14L:10D light cycle (Westerfield, 2007). The strains used in this study were AB (as wild type, WT), *kita(b5)*, referred to as *kita(lf)* (Parichy et al., 1999b), and *kitlga(tc244b)*, referred to as *kitlga(lf)* (Hultman et al., 2007). Previously published strains used include: *Tg(aox5:PALM-eGFP)* (Eom *et al*., 2015), *Tg(mitfa:BRAFV600E)* (REF Ceol 2011). Construction of new strains generated for this study is detailed below.

### DNA constructs

DNA constructs were built using the Gateway system (Life Technologies). Previously published entry clones used in this research are: pENTRP4P1r-Pmitfa (Ceol et al., 2011), pENTR-ERKKTRClover (Mayr *et al*., 2018), and p3E-polyA (Kwan et al., 2007). Previously published destination vectors include: pDestTol2pA2 ((Kwan *et al*., 2007). Using the entry clones described above, the following constructs were built with standard multisite Gateway reactions: pDestTol2pA2-Pmitfa:ERKKTRClover:pA, pDestTol2pA2-Pmitfa:nlsEGFP:pA, and pDestTol2pA2-Pmitfa:nlsmCherry:pA.

### Microinjection and transgenic fish

For transposon-mediated integration, 25 picograms of a construct was injected with 25 picograms of Tol2 transposase mRNA into zebrafish embryos at the single-cell stage. For injection into an existing Tol2-generated line, constructs were linearized and injected into single-cell embryos without transposase. To create the *Tg(mitfa:nlsEGFP)* transgenic line, pDestTol2pA2-Pmitfa:nlsEGFP:pA was injected into wild-type embryos, EGFP-positive larvae selected and the resulting adults outcrossed to wild-type animals. EGFP-positive larvae from these crosses were further outcrossed to wild-type animals to establish a stable *Tg(mitfa:nlsEGFP)* transgenic line.

### Melanocyte Destruction and Lineage Analysis

Adult zebrafish were treated with 750nM neocuproine in a beaker for 24 hours and then kept in individual tanks filled with fish water after drug washout. Prior to imaging fish were treated with 1mg/ml (-)-Epinephrine (+)-bitartrate salt (Acros Oragnics) for 2 minutes and then anesthetized with 0.17 mg/ml tricaine. Fish where then placed on their sides in a plastic Petri dish and imaged in the same locations using anatomical and cellular landmarks. Fish were viewed with a Leica DM550B microscope and images were captured with a Leica DFC365FX camera. The fish were then placed in fish water for recovery.

### Imaging and Quantitative Analysis

Brightfield images were adjusted for color balance, contrast, and brightness for clarity. Cells were counted along the flank’s middle melanocyte stripe (2^nd^ stripe down from the dorsum) under 20X magnification using FIJI ImageJ. Fractional regeneration was calculated as a ratio of the number of melanocytes within the stripe region at the specified time point relative to the same region prior to melanocyte destruction via neocuproine. Student’s t-tests were performed using GraphPad Prism 9. Lineage tracing was performed by aligning daily brightfield and fluorescent images using anatomical and cellular landmarks. *mitfa:nlsEGFP*-expressing cells were traced from Day 0 (prior to melanocyte ablation) until Day 10. Differentiation was recognized by melanization, and self-renewal was recognized by a division in which neither daughter cell differentiated.

### RT-PCR

Adult WT fish were dissected, skin tissue was placed in TRIzol (Thermo Fisher) and mechanically dissociated before RNA isolation and purification using the RNeasy kit (Qiagen) per manufacturer’s protocol. Full-length cDNA was synthesized with SuperScript III First Strand Synthesis kit (Thermo Fisher). Reaction mixes were comprised of SYBR Green RT-PCR master mix (Thermo Fisher), primers, and 25 ng cDNA (Supplementary Table 2). Analysis was performed using a StepOnePlus Real Time PCR System (Applied Biosystems). Samples were normalized to *β-actin* for loading control and *zfk8* for a surgical sampling control. Fold changes were calculated using ΔΔCt in Microsoft Excel.

### ERKKTR

To quantify ERK signaling, mClover intensity was measured in the nucleus and cytoplasm using Fiji’s measure intensity tool. Nuclear localization was defined by nlsmCherry expression. The ratio of cytoplasmic to nuclear signal intensity was calculated using Microsoft Excel. McSCs were randomly selected and quantified at timepoints before, during and after melanocyte regeneration.

### *aox5* intensity of McSC fates

McSCs were traced during regeneration and assayed for cell fate. Differentiation was recognized by melanization, and self-renewal was recognized by a division in which neither daughter cell differentiated. The *aox5* fluorescence intensity of these cells was then measured using Fiji. *aox5:PALM-eGFP* intensity was calculated in ImageJ by determining the mean intensity within a cell, subtracting the background, and log transforming the signal. Differences in *aox5* intensity between traced fates was visualized in Prism 9.

### scRNAseq sample prep and FACS enrichment

Fish skin tissue was enzymatically dissociated with Liberase TM (Sigma-Aldrich LIBTM-RO, 0.25mg/ml in dPBS) at 32°C for 20 minutes followed by manual titration with a glass pipette for 2 minutes. Cell suspensions were then filtered through a 70µm Nylon cell strainer. This single cell suspension was then spun down for 5 minutes at 1500pm in an Eppendorf 5810 R swing bucket centrifuge and resuspended in 0.1% BSA/5%FBS in dPBS for FACS enrichment.

Singlets were then isolated using a series of FSC-A vs SSC-A and FSC-A vs FSC-H gates. We then enriched for our *mitfa:nlsEGFP*-expressing cells using unlabeled WT animals as a negative control. All samples were kept on ice in 0.1%BSA/5%FBS except during enzymatic dissociation. All surfaces encountering the cell solution were coated in 1%BSA before use.

### scRNAseq regeneration sampling, library construction sequencing

For each time point before (day 0) and during (days 1,2,3,5 and 10) regeneration we captured transcriptomes from WT and *kita(lf)* fish. To generate sufficient material for single-cell capture 30 zebrafish lateral skins were dissected for each time point and an estimated 10,000 EGFP-positive cells per sample were obtained via FACS. For each sample we targeted 4,000-6,000 cells for capture using the 10X Genomics Chromium platform with one sample per lane.

Libraries were prepared using the Single Cell 3’ kit (v3.1). Quality control and quantification assays were performed using a Qubit fluorometer (Thermo Fisher) and a fragment analyzer (Agilent). Libraries were sequenced on an Illumina NextSeq 500 using 75-cycle, high output kits with read 1: 28 cycles, index: eight cycles, read 2: 50 cycles. Each sample was sequenced to an average depth of 216 million total reads resulting in an average read depth of approximately 59,000 reads/cell.

### scRNAseq sequencing analysis

We ran the below pipelines in the DolphinNext environment (https://github.com/UMMS-Biocore/dolphinnext) using the Massachusetts Green high Performance Computing Cluster (Yukselen et al., 2020). Raw base call (BCL) files were analyzed using CellRanger (version 3.1.0). Cell Ranger “mkfastq” was used to generate FASTQ files and “count” was used to generate raw feature-barcode matrices aligned to the Lawson Lab zebrafish transcriptome annotation v4.3 (Lawson et al., 2020). Cell Ranger defaults for selecting cell-associated barcodes versus background empty droplet barcodes were used.

### scRNAseq downstream data analysis

Filtering retained cells that expressed between 200 and 6000 unique genes, and a mitochondrial fraction less than 10 percent. These parameters resulted in retention of 29,453 WT cells and 24,724 *kita(lf)* cells across 14 total samples. Each dataset was further pre-processed by log normalizing each dataset with a scale factor of 10,000 and finding 2,000 variable features using the default ‘vst’ method in seruat3.0 (Stuart *et al*., 2019). Datasets were integrated using the WT datasets as references in the “FindAnchors” step (Stuart *et al*., 2019). After integration the integrated dataset was scaled to regress out differences driven by inequalities in total RNA count and mitochondrial capture. We then reduced computed 100 principal components on 2000 variable features. The top 60 principal components were used for construction of a UMAP and neighbor finding. During clustering we utilized a resolution parameter of 1.0.The resulting 33 Louvian clusters were visualized in 2D and/or 3D space and were annotated using known biological cell-type markers. Visualization of the UMAP and calculation of DEGs split by sample revealed no observable strong batch effects. Changing any of the above parameters yielded similar cell type identifications and clustering structures. Both 2000 or 4000 variable features revealed similar results in final cell clustering and DEG analysis. Similarly, using between 20 and 80 principal components and a resolution parameter between 0.5 and 1.2 revealed similar UMAP visualizations and clustering, underlining the robustness of the approach.

### NICHENETR

Our analysis identifying potential ligand/receptor pairs regulating McSC behavior during regeneration utilizes the open-source R implementation of NicheNetR available at GitHub (github.com/saeyslab/nichenetr). We followed the default parameters for average logFC and percent expression. For elucidating ligand/receptor interactions we assigned all non-McSC cell populations as defined by clustering in Seurat as potential “sender cells” and McSCs as “receiver cells”. We then filtered down receptor ligand pairs to the “Bona-Fide” literature supported interactions using NicheNetR’s built in lists.

### Trajectory analysis and pseudotime

Graph trajectory learning as well as pseudotime was calculated using default parameters on a UMAP visualization of McSCs and melanocytes obtained in Seurat and imported into monocle3 (Saunders *et al*., 2019). To normalize for differences in real-time sampling, cells were randomly downsampled to the lowest common denominator so that real-time sample size differences would not impact pseudotime distribution calculation. RNA velocity splicing mechanics were calculated using default parameters in velocyto (La Manno *et al*., 2018). Then velocity embeddings and streams were calculated on the Seurat derived UMAP embedding using default parameters in scvelo (Bergen *et al*., 2020).

### Marker Calculation and Comparison to Murine Datasets

Murine sample data were obtained from GEO GSE147299. Data were processed in Seurat using methods described above. Cells from *BMP*-KO samples as well as non-melanocyte lineage keratinocytes were then removed. Signature expression was calculated used “FindMarkers” in each dataset. Markers were then converted to murine orthologs using HGNC orthology tables as a key. Zebrafish signature scores were computed for murine data using Seurat’s “AddModuleScore.” Scores between cell types were visualized using a DotPlot..

### Comparisons to fibroblast and keratinocyte data

Murine sample data of fibroblasts and keratinocytes were obtained from GEO GSE129218. Data were processed in Seurat using methods described above. Gene expression profiles for canonical fibroblast and keratinocyte marker genes were visualized alongside *Kitlg* expression using Feature plots.

### Statistics

Student’s t-tests were performed using GraphPad Prism 9. Wilcoxon rank-sum tests were calculated in R[version 4.0.0] (RCoreTeam, 2020). Cluster population shifts analyzed using differential proportion analysis (DPA) using default parameters (Farbehi *et al*., 2019).

### Data availability

Raw Data and 10X output files are deposited in GSE190115.

---

**Supplemental Figure 1.**
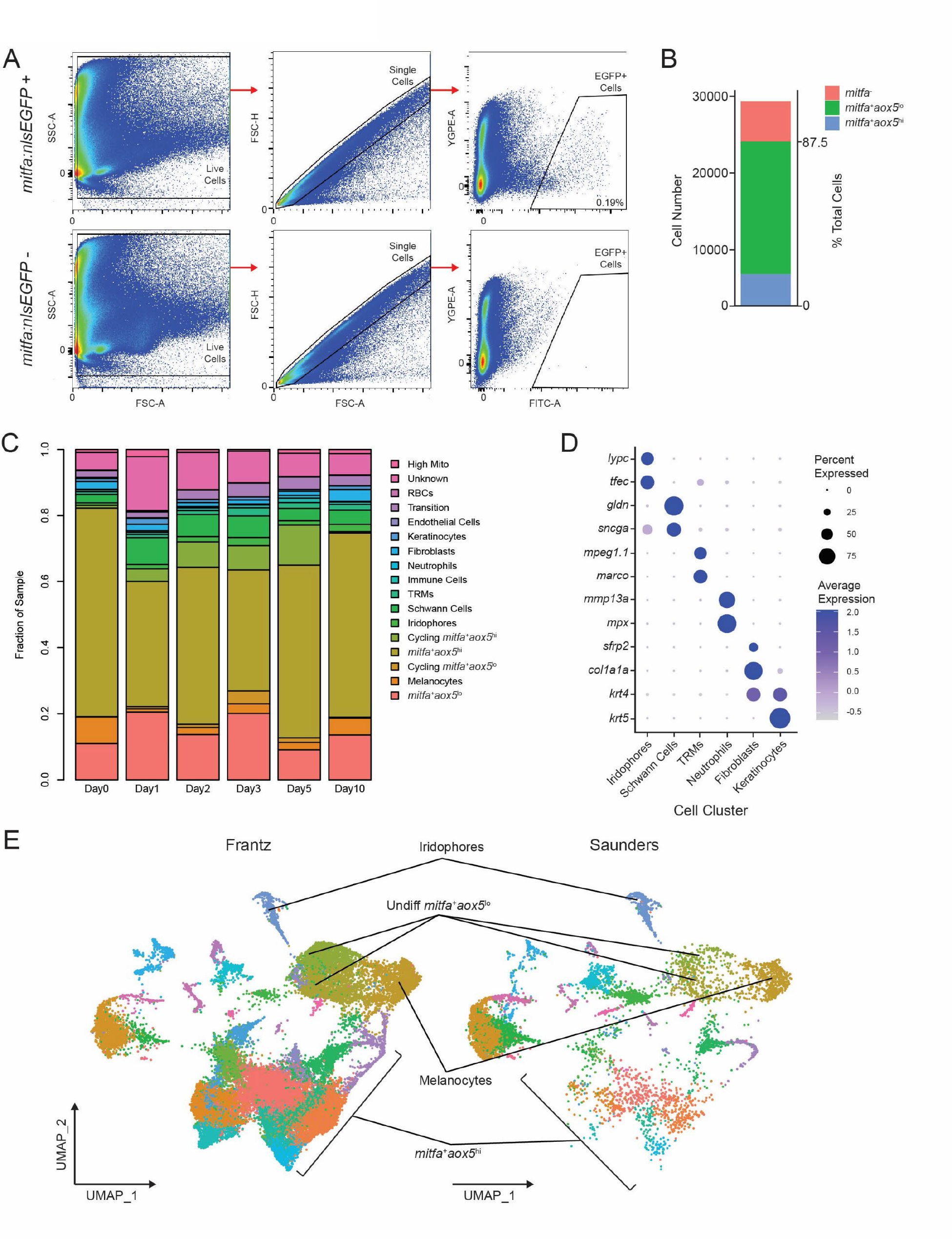
Isolation strategy and characteristics of *mitfa:nlsEGFP*-expressing cells. (A) Representative FACS plots and gating strategy to isolate *mitfa:nlsEGFP*-positive cells from zebrafish skin. Top, isolation from skin of *Tg(mitfa:nlsEGFP)* zebrafish, and, bottom, from negative control skin of wild-type zebrafish. (B) Percentage of *mitfa*-expressing cells from FACS isolation, based on scRNAseq analyses. (C) Proportions of cell types across all *Tg(mitfa:nlsEGFP)* samples. High Mito: high mitochondrial read fraction cells; Unknown: cells with unclear literature supported markers; RBCs: red blood cells; Transition: cells spanning a geographic transition between the *mitfa*-expression populations; TRMs: tissue-resident macrophages. (D) Expression of cell markers for cell clusters shown in Figure 1A. Dot sizes represent percentage of cells in the cluster expressing the marker and coloring represents average expression. (E) Comparison of our *mitfa:nlsEGFP*-enriched dataset from zebrafish adults with *sox10:Cre:mCherry*-enriched dataset from larval zebrafish (Saunders *et al*., 2019).

**Supplemental Figure 2.**
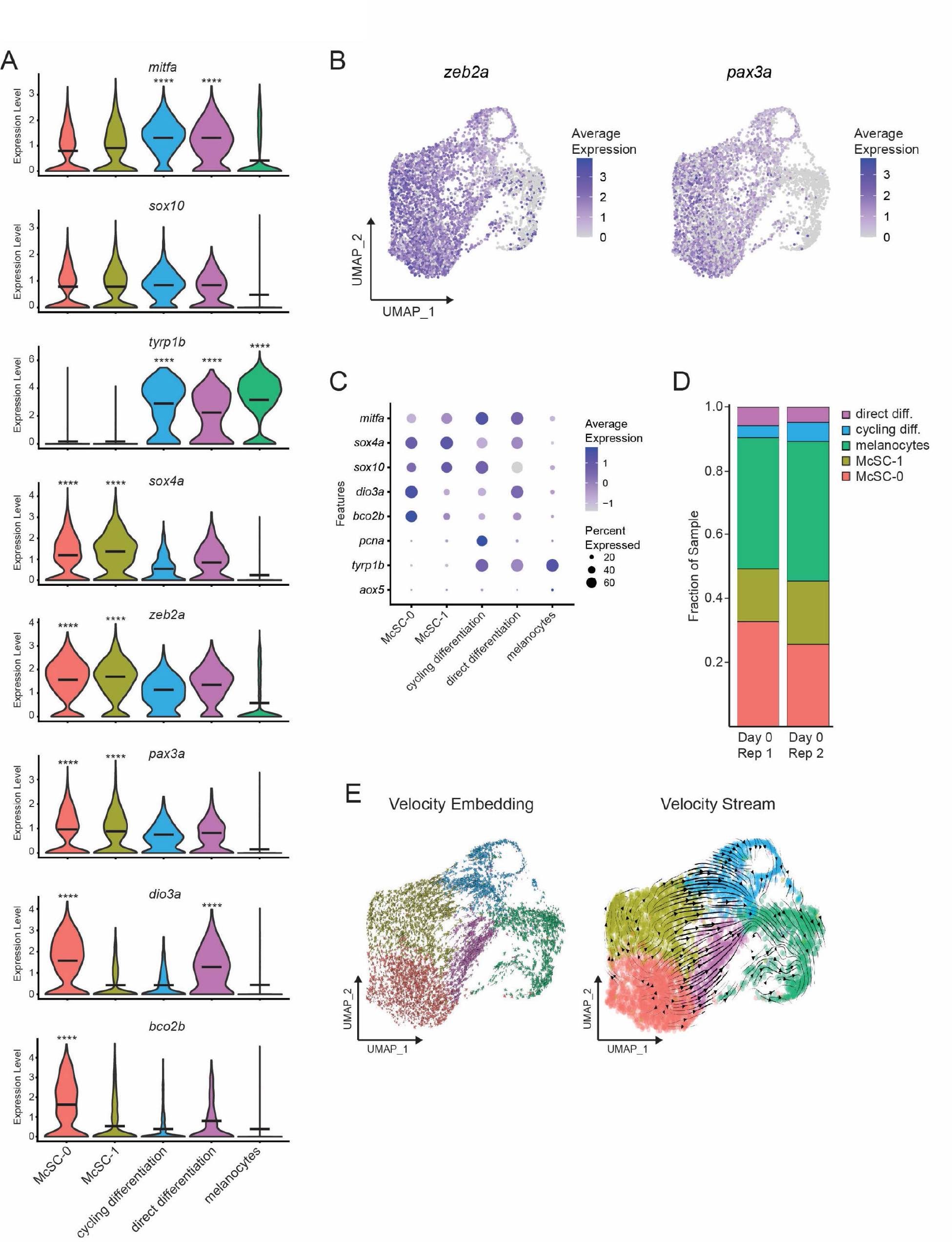
Single-cell profiling supports McSC identity and subpopulation dynamics. (A) Violin plots of cell subpopulation expression of *mitfa*, pigment cell progenitor (*sox10*), melanin biosynthesis (*tyrp1b*), stem cell (*sox4a*), and neural crest (*zeb2a*, *pax3a*) genes. Genes distinguishing the two McSC clusters (*dio3a*, *bco2b*) are also visualized. Black bars represent mean gene expression. P values calculated by Wilcoxon rank sum test, **** p < 0.0001 (B) Expression of neural crest genes *zeb2a* and *pax3a* and their enrichment in McSC-0 and McSC-1 subpopulations as feature plots on the UMAP plot from Figure 2A. (C) Dot plot demonstrating gene expression differences between *mitfa*^+^*aox5*^lo^ cell subpopulations. Dot sizes represent percentage of cells in the cluster expressing the marker and coloring represents average expression. (D) Comparison of *mitfa^+^aox5*^lo^ cell subpopulation proportions between different scRNAseq samples of unperturbed (day 0) wild-type zebrafish skin. (E) RNA splicing mechanics scored using velocity embedding (velocyto; (La Manno *et al*., 2018) and visualized through velocity stream analysis (scvelo; (Bergen *et al*., 2020)) on the UMAP plot from Figure 2A.

**Supplemental Figure 3.**
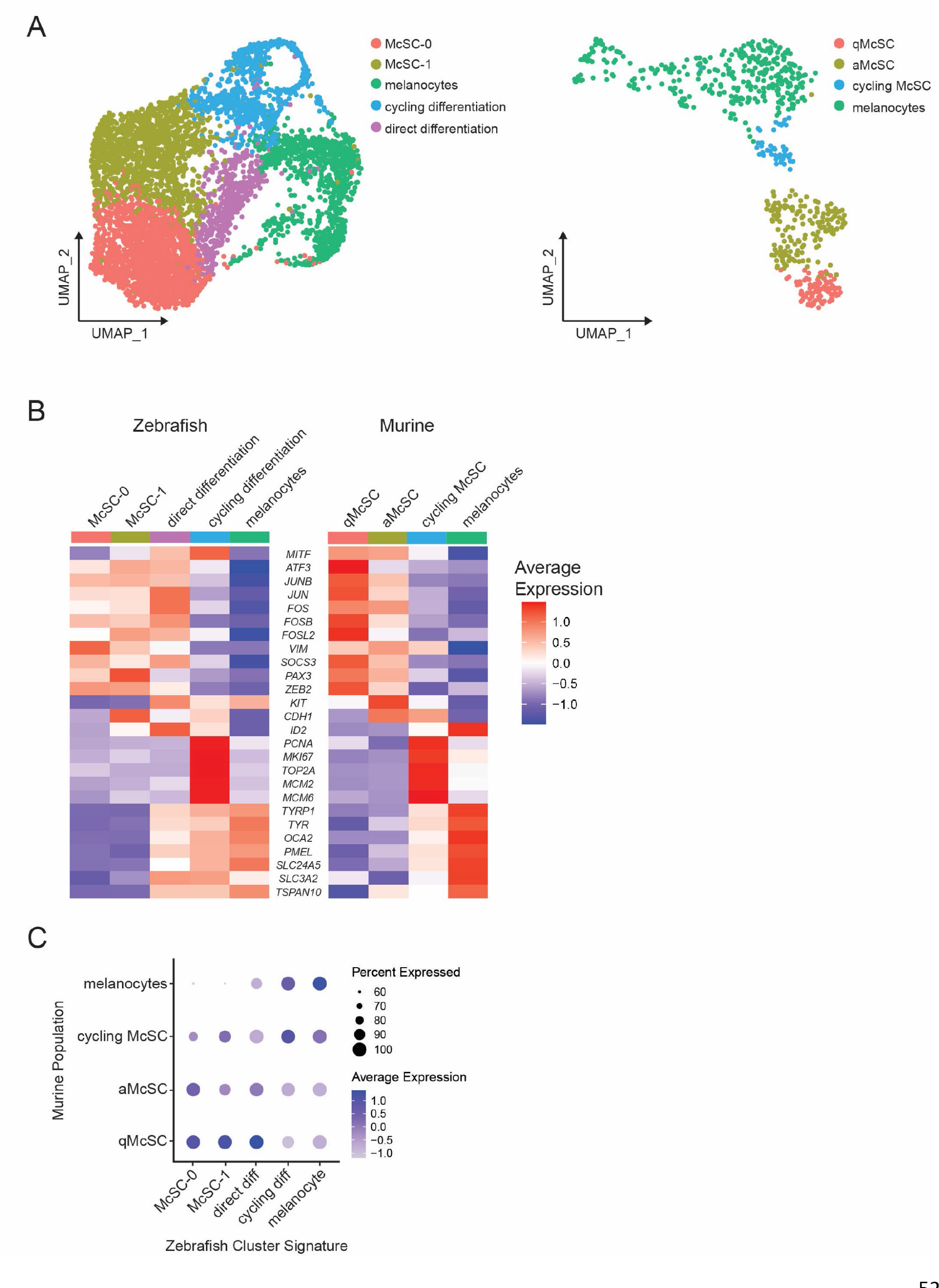
Zebrafish McSC regenerative signature is conserved across species. (A) Left, UMAP subclustering of *mitfa^+^aox5*^lo^ cells before, during, and after melanocyte regeneration as in Figure 2A (n=5,619 cells). Right, integrated sub-clustering of murine McSCs, intermediate cell populations and melanocytes during hair cycle turnover (n=626 cells); qMcSCs, quiescent McSCs; aMcSCs, activated McSCs (Infarinato *et al*., 2020). (B) Heatmaps of melanocyte-lineage cluster markers conserved across zebrafish and murine scRNAseq datasets. Gene labels are human orthologs of genes expressed as cluster markers in both zebrafish (left) and murine (right). Gene expression is row normalized. (C) Dot plot visualizing conserved regenerative transcriptional states across species. Zebrafish subpopulation signatures were calculated using the FindMarkers feature in Seurat (Satija et al., 2015; Stuart *et al*., 2019) from zebrafish subpopulations then these zebrafish signatures were plotted onto murine subpopulations (Infarinato *et al*., 2020). Dot sizes represent percentage of cells in the murine subpopulation expressing the zebrafish gene signature and coloring represents average expression.

**Supplemental Figure 4.**
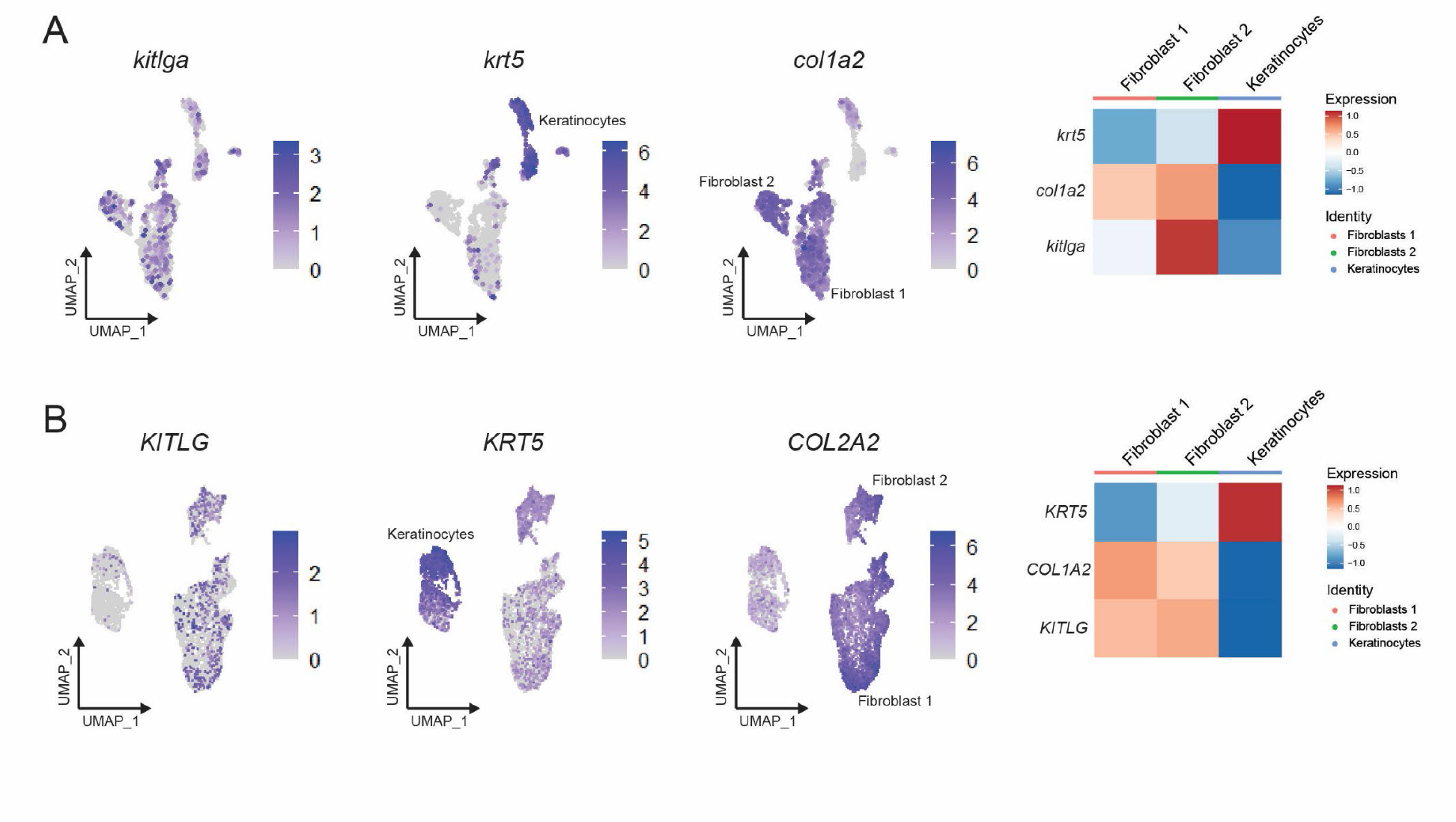
*kitlga* ligand expression in zebrafish keratinocytes and fibroblasts is similar to that observed in human cells. (A) Left, feature plots of zebrafish *kitlga* expression, keratinocyte marker *krt5*, and fibroblast marker *col1a2*. Right, heatmap plots of *kitlga*, *krt5* and *col1a2* expression in keratinocytes and fibroblasts. (B) Left, feature plots of human *KITLG* expression, keratinocyte marker *KRT5*, and fibroblast marker *COL2A2* (Joost *et al*., 2020). Right, heatmap plots of *KITLG*, *KRT5* and *COL2A2* expression in keratinocytes and fibroblasts.

**Supplemental Figure 5.**
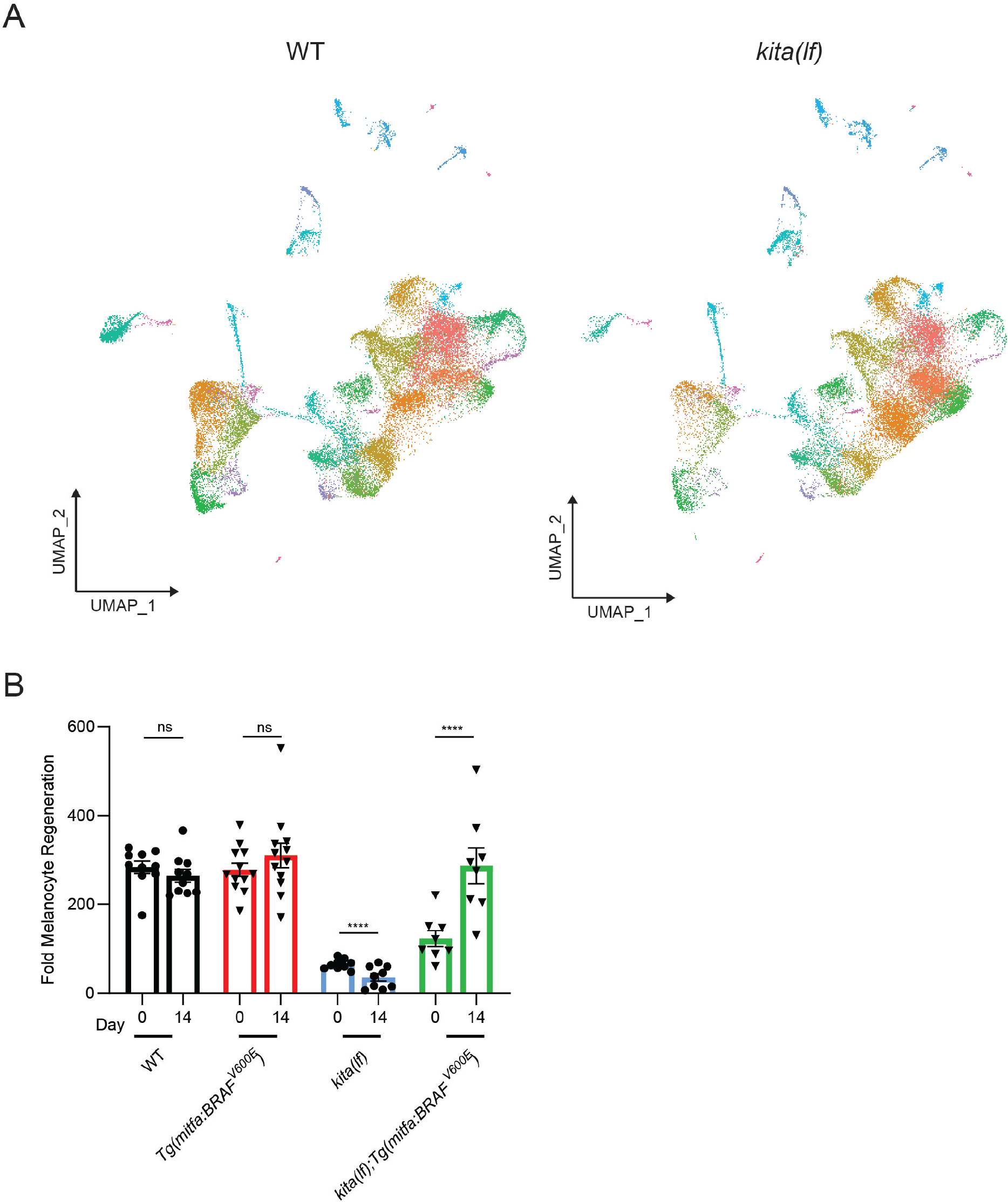
*kita(lf)* animals with constitutively active BRAF regenerate a similar number of melanocytes as wild-type animals. (A) Unsupervised clustering of all wild-type (left, n=29,453 cells) and *kita(lf)* (right, n=24,724 cells) cells profiled by scRNAseq. (B) Mean number of melanocytes from the middle stripe per field. + SEM is shown; wild type n = 10, *Tg(mitfa:BRAFV600E)* = 12, *kita(lf)* = 9, *kita(lf); Tg(mitfa:BRAFV600E)* = 8 fish. P values calculated by Student’s t-test, **** p < 0.0001; ns, not significant.

**Supplemental Figure 6.**
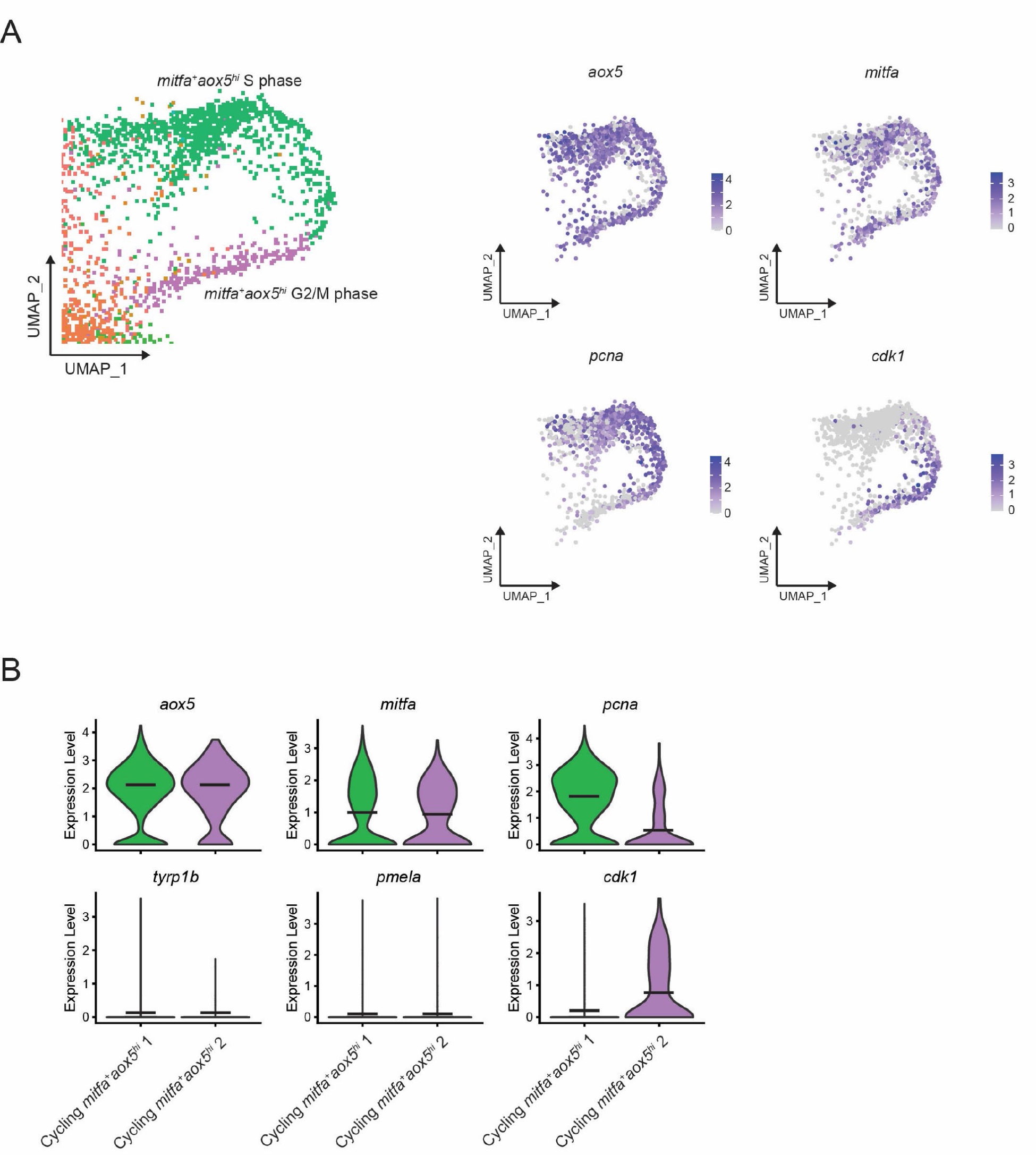
Gene expression in *mitfa*^+^*aox5*^hi^ cycling subpopulations. (A) Left, enlargement of the *mitfa*^+^*aox5*^hi^ cycling subpopulation from Figure 6A showing two clusters. Right, feature plots of expression of pigment cell markers and cell cycle genes in the *mitfa*^+^*aox5*^hi^ cycling clusters. (B) Violin plots of gene expression of pigment cell markers and cell cycle genes of the clusters seen in Supplementary Figure 2A. Black bars represent mean gene expression.

**Supplemental Figure 7.**
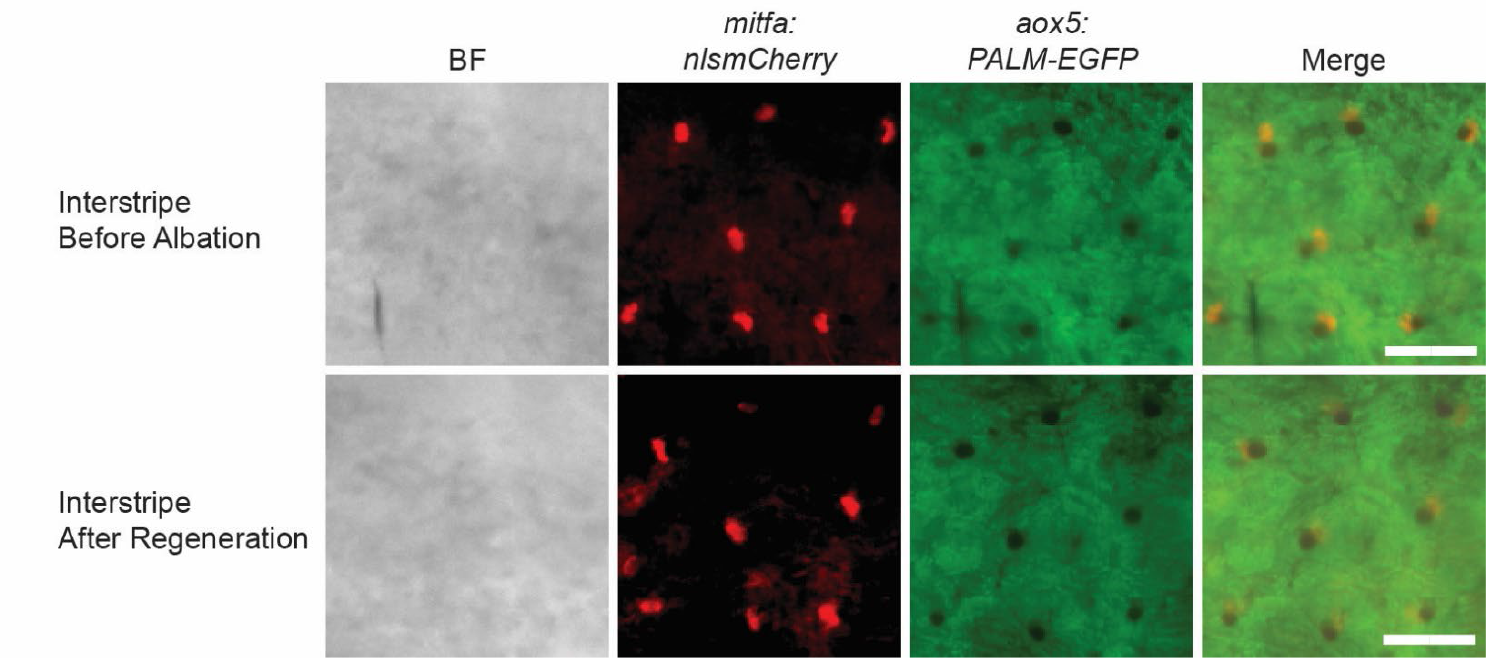
Lack of interstripe xanthophore divisions following neocuproine treatment. Representative images from lineage tracing of interstripe xanthophores from *Tg(mitfa:nlsmCherry); Tg(aox5:PALM-EGFP)* animals following neocuproine treatment. Scale bar = 60µm.

## Notes

### Competing Interest Statement

The authors have declared no competing interest.

